# Estimating the cumulative impact and zone of influence of anthropogenic features on biodiversity

**DOI:** 10.1101/2022.06.14.495994

**Authors:** Bernardo Brandão Niebuhr, Bram Van Moorter, Audun Stien, Torkild Tveraa, Olav Strand, Knut Langeland, Per Sandström, Moudud Alam, Anna Skarin, Manuela Panzacchi

## Abstract

1. The concept of cumulative impacts is widespread in policy documents, regulations, and ecological studies, but quantification methods are still evolving. Infrastructure development usually takes place in landscapes with preexisting anthropogenic features. Typically, their impact is determined by computing the distance to the nearest feature only, thus ignoring the potential cumulative impacts of multiple features. We propose the *cumulative ZOI approach* to assess whether and to what extent anthropogenic features lead to cumulative impacts.
2. The approach estimates both effect size and zone of influence (ZOI) of anthropogenic features and allows for estimation of cumulative effects of multiple features distributed in the landscape. First, we use simulations and an empirical study to understand under which circumstances cumulative impacts arise. Second, we demonstrate the approach by estimating the cumulative impacts of tourist infrastructure in Norway on the habitat of wild reindeer (*Rangifer t. tarandus*), a nearly-threatened species highly sensitive to anthropogenic disturbance.
3. Simulations show that analyses based on the nearest feature and our cumulative approach are indistinguishable in two extreme cases: when features are few and scattered and their ZOI is small, and when features are clustered and their ZOI is large. Empirical analyses revealed cumulative impacts of private cabins and tourist resorts on reindeer, extending up to 10 and 20 km, with different decaying functions. Although the impact of an isolated private cabin was negligible, the cumulative impact of ‘cabin villages’ could be much larger than that of a single large tourist resort. Focusing on the nearest feature only underestimates the impact of ‘cabin villages’ on reindeer.
4. The suggested approach allows us to quantify the magnitude and spatial extent of cumulative impacts of point, linear, and polygon features in a computationally efficient and flexible way and is implemented in the `oneimpact` R package. The formal framework offers the possibility to avoid widespread underestimations of anthropogenic impacts in ecological and impact assessment studies and can be applied to a wide range of spatial response variables, including habitat selection, population abundance, species richness and diversity, community dynamics, and other ecological processes.

## 1 Introduction

Land use change and infrastructure development are increasing at an accelerating pace worldwide (Ibisch *et al*., 2016; Venter *et al*., 2016), including all global biodiversity hotspots (Hu *et al*., 2021), and are among the main causes of an unprecedented biodiversity decline (Beńıtez-López *et al*., 2010; Newbold *et al*., 2015, IPBES, 2019). Most infrastructure development takes place in areas already affected by multiple sources of disturbance (Barber *et al*., 2014) and, therefore, anthropogenic features are often clustered in the landscape. Understanding biodiversity responses to spatially co-occurring features is crucial to adequately assess their total impact. Indeed, the impact of new anthropogenic features might add and spatially interact to that of preexisting ones, leading to cumulative impacts larger than that of single features in isolation (Box 1; Johnson & St-Laurent, 2011). Adequately quantifying anthropogenic cumulative impacts is crucial to promote ecological sustainability in land planning, to prevent habitat loss, and to inform robust mitigation and offset measures (Gillingham *et al*., 2016; Laurance & Arrea, 2017). Most environmental impact assessment studies focus on single infrastructure projects at small spatio-temporal scales (Johnson, 2011), and even broad-scale ecological studies typically consider only the impact of the nearest anthropogenic feature, thus ignoring cumulative impacts of multiple co-occurring features (e.g. Torres *et al*., 2016). This however relies on the strong assumption that the impact is caused only by the anthropogenic feature closest to an species’ location, and co-occurring features have no additional impact. Although there have been efforts to better define, review, and outline cumulative impacts (Gillingham *et al*., 2016; Johnson & St-Laurent, 2011), we still lack a comprehensive theory and framework to understand and quantify cumulative impacts, and thus concretely help sustainable land use planning. This paper aims to take this process one step further by proposing the *cumulative ZOI approach* to quantify the impact of multiple anthropogenic features on species, communities, and ecological processes.

### Box 1

#### Definitions

**Impact** We use the term to describe the consequences of infrastructure, land use, human disturbance, and any spatial feature on a biological response variable, such as species’ occurrence, biodiversity, or ecological processes. Therefore, impacts represent the functional responses of species and processes to human activity (Johnson & St-Laurent, 2011). We analytically decompose the impact *I* into its **effect size** *β* and its spatial component, the **zone of influence (ZOI)** *ϕ*, so that *I* = *β · ϕ*. A given anthropogenic feature (e.g. tourist cabin) might affect a certain process (e.g. species occurrence) strongly or weakly (*β*), and this impact might decrease rapidly with distance or extend over several kilometers (ZOI, *ϕ*).

**Cumulative impacts** can result from the interaction between multiple features of a given type – our focus here –, from the impact of different types of infrastructure (e.g. houses, turbines, roads, or dams) or from top-down or bottom-up ecological cascades. Cumulative impacts of multiple features depend on the number of features, their spatial distribution, configuration, and co-occurrence with other disturbance types, and might differ across species or processes, possibly leading to stronger impacts (negative or positive), if compared to the impact of a single isolated feature.

**Effect sizes** express how strongly a given biological response is affected by a type of disturbance at the point in space where the disturbance is located. Here, the effect size is given by the estimated model coefficients *β* (eq. 2).

**Zone of Influence** The **ZOI** represents the function *ϕ* defining how the effect size of an anthropogenic feature changes with the distance, that is, it represents how the impact spreads throughout space. The ZOI might be any function *ϕ*(*d, r*) that assumes value 1 at the origin and decreases towards zero as the distance *d* from the feature increases. The ZOI is defined by its **shape** and **radius *r***. The ZOI **shape** determines how *ϕ* decreases with distance, or whether it remains constant up to a threshold distance *r* (see Fig. 1A and Appendix A for examples). The ZOI **radius** *r* is the maximum distance from the feature where it affects a given biological response. For some shapes (e.g. threshold, linear decay) *r* is the distance at which *ϕ* reaches zero (Fig. 1A). For non-vanishing functions (e.g. Gaussian decay), a cutoff must be set to define *r* – e.g. the minimal distance where *ϕ* is below 0.05. While in landscape ecology the ZOI radius is often called the *scale of effect* (e.g. Moraga *et al*., 2019), we refer to the ZOI radius to avoid confusion from different definitions of *scale*.

**ZOI metrics** When multiple features of an infrastructure are present, we can compute two ZOI metrics: the ZOI of the nearest feature only (*ϕ_nearest_*, eq. 4) and the cumulative ZOI of multiple features (*ϕ_cumulative_*, eq. 5), which is the sum of the ZOI of each feature (Fig. 1B). Each of these metrics is a different predictor, and the estimated ZOI is determined through statistical fitting and model selection.

Anthropogenic features directly affect species in the area where they are physically present (e.g. through habitat loss or road kills), but their effect might also extend far beyond the features themselves, for instance by causing avoidance responses and reducing the probability of animal occurrence in their proximity (Johnson *et al*., 2005; Torres *et al*., 2016). On a broader landscape perspective, this can lead to the obstruction of movement or migration corridors, which in turn can prevent access to functional areas further away, with far reaching consequences for species distribution and population dynamics (Panzacchi *et al*., 2016; Dorber *et al*., 2023; Van Moorter *et al*., 2021, 2023, in press). Therefore, two intrinsically related dimensions must be estimated in cumulative impact studies: the effect size of the impact and the size of the area affected (Box 1; Johnson & St-Laurent, 2011). The *effect size* indicates how strongly a feature influences the focal species or process, and it is generally estimated through a combination of biological and environmental data through statistical modeling (Box 1; Polfus *et al*., 2011). The *zone of influence* (ZOI) defines the area within which the impact of the feature is detectable, is commonly expressed using the radius of a circle with the feature in the origin, and delimits the area affected (Box 1; Boulanger *et al*., 2021; Polfus *et al*., 2011).

The impact of co-occurring spatial features can accumulate over space (and time), as a linear or non-linear function of the impact of each individual feature. Such cumulative impacts are commonly appraised by reclassifying the features into larger units; for instance several point features representing buildings may be reclassified as a polygon representing an urban area, or several wind turbines as a wind park (Torres *et al*., 2016). For determining the ZOI, two approaches are typically used: either measuring the distance to the features or their density. The first framework focuses on the concept of ecological thresholds (Ficetola & Denöel, 2009) and often estimates the ZOI by modelling the species’ response as a function of distance from disturbance using piecewise regression or other regression models (e.g. exponential decay or generalized additive models; Ficetola & Denöel, 2009; Skarin *et al*., 2018). This approach typically considers only the distance to the nearest feature and assesses ZOI thresholds only for one or a few types of anthropogenic feature (e.g. Boulanger *et al*., 2021), since its computation requires repeated fitting and becomes impractical in a broader context (Lee *et al*., 2020).

The second approach estimates ZOI focusing on the spatial and temporal *scales of effect* of the species-habitat relationships (e.g. Zeller *et al*., 2017). In this context, the number of features is averaged at several spatial extents (Moraga *et al*., 2019; Laforge *et al*., 2015), creating a series of disturbance density maps (McGarigal *et al*., 2016). Each of these maps is tested against a biological response variable to assess the spatial scale at which the relationship is strongest, commonly through measures of model performance and explanatory power (such as R^2^, AIC, or BIC, or through the fitted model coefficients; Huais 2018). Multi-scale analyses brought important advances into spatial ecology and environmental impact studies (e.g. McGarigal *et al*., 2016). However, these studies are rarely used in the context of cumulative impact assessments (but see Polfus *et al*., 2011).

Here, we propose the *cumulative ZOI approach* to detect the occurrence of cumulative impacts on biological variables and to quantify them assuming additive effects of multiple features. In the approach, the zone of influence (ZOI) describes how the impact of a feature decreases with distance from the feature, and we use a model selection approach to determine a suitable functional form of the ZOI. The approach allows estimating the effect of both the nearest feature only, and of the cumulative impact of multiple features (Box 1, Fig. 1). For simplicity, in this paper we focus on the impact of features of the same type, although the approach can be extended to different feature types. First, we perform simulations to illustrate the performance of each of these two metrics in explaining species’ responses to the same anthropogenic features under different spatial configurations i.e. scattered vs. clustered. Second, we demonstrate the approach by assessing the cumulative impact of private cabins and tourist resorts in Norway on the habitat selection of the tundra’s flagship species, reindeer. We developed the `oneimpact` R package to allow implementation of the approach in R (R Core Team, 2020) and GRASS GIS (GRASS Development Team, 2017). Given the occurrence of multiple anthropogenic features, the cumulative ZOI approach allows to: (i) evaluate whether there is evidence for cumulative impacts, or whether the impact of the nearest feature is sufficient to capture the species’ spatial response; (ii) quantify the cumulative impact; and (iii) estimate the ZOI and spatial decay function for multiple types of features. Although the approach was developed for infrastructure and anthropogenic disturbance factors, it can be used for any spatial predictor, including landscape and natural variables, and can be extended to many types of ecological responses, such as animal movement, species occurrence and abundance, and population fecundity and genetics (Panzacchi *et al*., 2016; Moraga *et al*., 2019; Collevatti *et al*., 2020).

**Figure 1:**
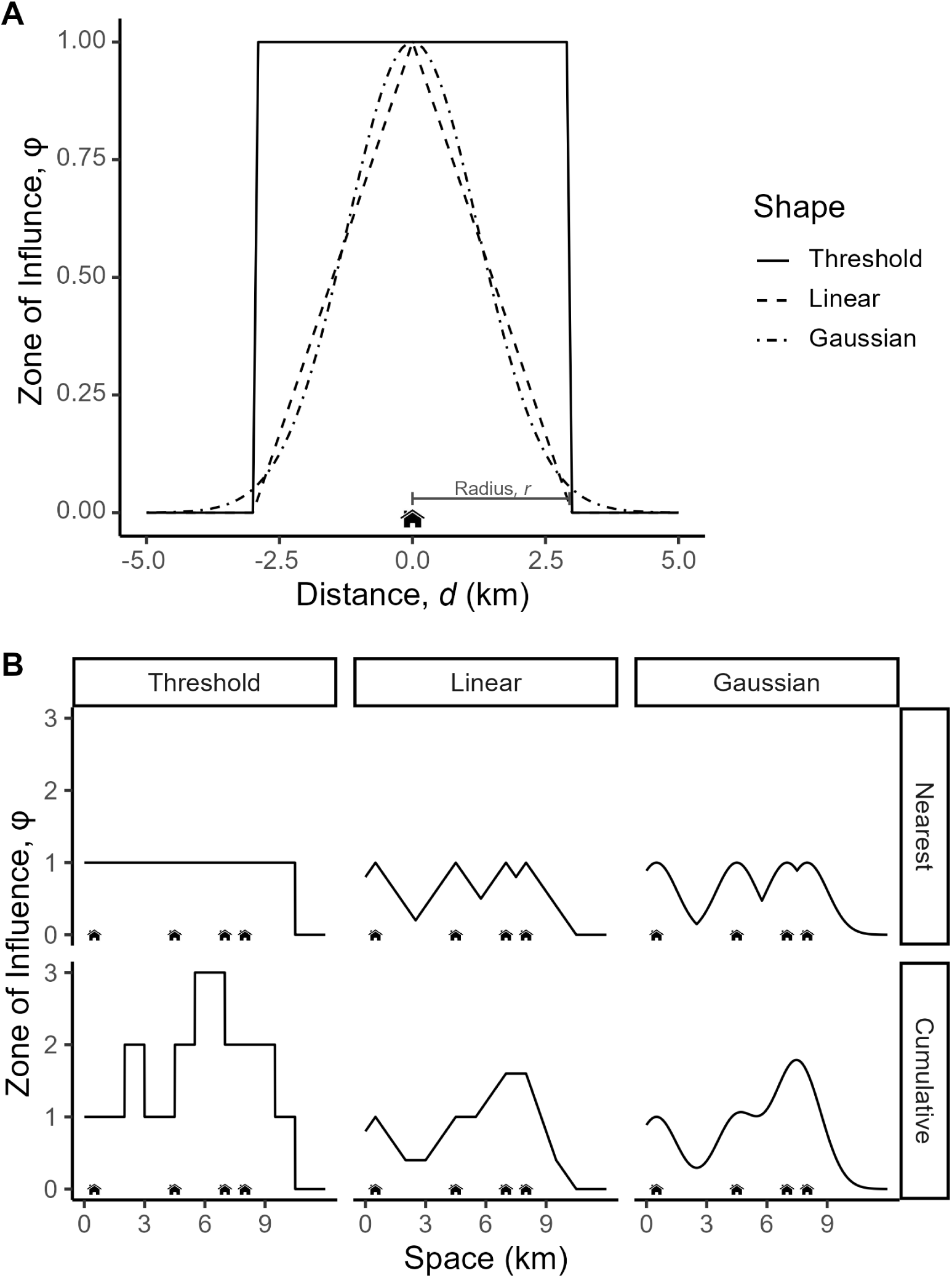
Illustration of the ZOI of an anthropogenic feature in one dimensional space, using houses as example. (A) Examples of ZOI functions *ϕ* with different shapes of decay with distance from feature, *d*. A house has only influence within its ZOI radius (here *r* = 3 km). For the threshold function, the influence remains constant within the ZOI and drops to zero beyond it, while for both the linear and the Gaussian functions it decreases monotonically for *d* ≤ *r*. (B) Representation of the ZOI of multiple houses considering only the nearest feature (*ϕ_nearest_*, upper row) or the cumulative ZOI of multiple features (*ϕ_cumulative_*, bottom row), for different shapes. If only the nearest house is considered, *ϕ_nearest_* does not exceed one; when all houses act cumulatively, *ϕ_cumulative_* can be higher than one.

## 2 Defining cumulative impact and ZOI for multiple features

Hereafter for simplicity we refer to *infrastructure* or *features* to refer to any spatial feature related to anthropogenic disturbance, including buildings (e.g. tourist areas), industrial areas (e.g. wind power), linear infrastructure (e.g. roads, hiking trails), land use practices, and human disturbance factors (e.g. tourist volume). To illustrate the approach, we also focus on infrastructure of the same type (e.g. cabins), although the approach can and should be extended to include different types of anthropogenic factors in an area. We first derive a metric to describe the impact of multiple anthropogenic features on a biological response variable. Despite the usefulness of the approach in different contexts (see Section 3), for illustration we use a habitat selection analysis, where the aim is to discriminate environmental conditions selected or avoided by animals based on ecological data (e.g. species’ occurrence or movements) and a use-availability design (Fieberg *et al*., 2021). The habitat selection function (HSF) *w*(**X**, **Z**) is proportional to the probability of selection of a given resource unit, estimated from the frequency of used vs. available resource units. The HSF *w*(**X**, **E**) is function of a matrix of spatial predictor variables describing infrastructure, **X**, for which we want to estimate the impact and ZOI, and a matrix of other environmental variables, **E** (e.g. temperature, vegetation, altitude, or topography). In its parametric form, the HSF may be represented by

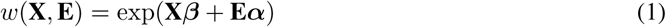

where ***β*** and ***α*** are vectors of coefficients for **X** and **E**. The first term can be written in vector form as

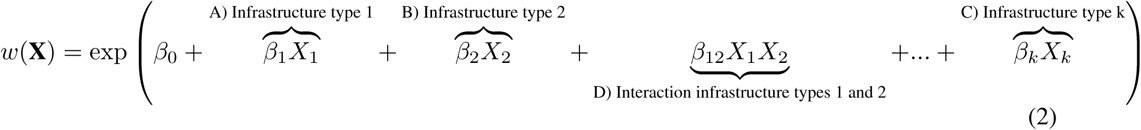

where we define each term *I_k_* = *β_k_X_k_* as the *impact* of a given anthropogenic feature of type k, and the impact is decomposed into its effect size *β_k_* and a spatial component *X_k_*. In terms of ecological interpretation, exp(*β_k_*) might be understood as the relative selection strength (Avgar *et al*., 2017), and exp(*I_k_*) as how this relative selection strength varies in space, for infrastructure type *k* (Fieberg *et al*., 2021).

In this formulation, *the cumulative impact of different types of infrastructure* is given by the additive impacts of the *k* infrastructure types (e.g. terms A, B, and C in equation 2) and possibly by interaction terms between variables (such as term D in equation 2, with an interaction coefficient *β*_12_), that allows for non-linear, joint effects caused by the co-occurrence of different types of infrastructure.

In our definition of impact, *β* and *X* are independent and *the cumulative impact of multiple features of the same type* is determined by the spatial component of the impact, *X*. We start by defining the ZOI as a function *ϕ*, a curve that represents how the infrastructure impact changes with distance (Box 1). The ZOI of each anthropogenic feature may follow different shapes: it may be either constant (threshold ZOI) or decrease with distance (e.g. linear and Gaussian ZOI, Fig. 1A). More broadly, *ϕ* = *f* (*d, r*) is any decay function that has a maximum value 1 where the feature is located and decreases toward zero as the Euclidean distance *d* increases, and possibly vanishes at a given point, the ZOI radius *r* (Box 1, Appendix A). Determining the ZOI shape and radius is an empirical problem (Miguet *et al*., 2017). The simplest assumption, widely used in the literature, is that all areas within the ZOI are affected equally (a buffer zone around features; e.g Panzacchi *et al*., 2013)). However, it is likely more reasonable to consider a higher *ϕ* closer to the disturbances (Skarin *et al*., 2018; Zeller *et al*., 2017). When multiple features coexist in the same area, we can define two ZOI metrics: the ZOI based on the nearest feature alone, *ϕ_nearest_*, and the cumulative ZOI of multiple features, *ϕ_cumulative_*(Box 1, Fig. 1B). For instance, for *ϕ_nearest_*the ZOI is assumed to be similar when approaching an isolated house and a small village, while for *ϕ_cumulative_* the ZOI of nearby houses adds up and will be greater than the ZOI of a single isolated house (Fig. 1B).

To translate these measures into a mathematical form, we can decompose each of the impact terms (i.e. A, B, C, …), in equation 2. Suppose that in the landscape there are *n_k_* features of type *k*, and let the ZOI of feature *i* of type *k* follow *ϕ_i,k_* = *f* (*d_i,k_*; *r_k_*), where *d_i,k_* is the distance to feature (*i_k_*) and *r_k_* is its ZOI radius. We can sum the effect of each feature so that the impact terms in equation 2 become:

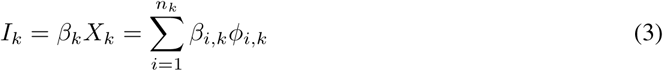

Typically, only the nearest feature is considered, which results in the implicit assumption that *β_i_* = 0 for all but the nearest feature. Thus, eq. 3 turns into:

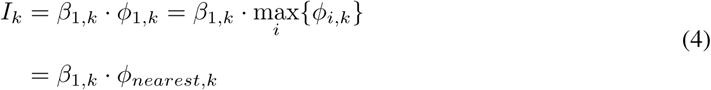

where *ϕ_nearest,k_* is the ZOI of the nearest feature (*i* = 1) of type *k* (see Fig. 1B). However, a possibly more reasonable assumption would be that *β_i,k_* = *β*_(_*_i_*_+1)_*_,k_* = *…* = *β_k_*, i.e. that all features of a given type have the same ZOI and all *β*’s are identical. Thus, eq. 3 is reduced to:

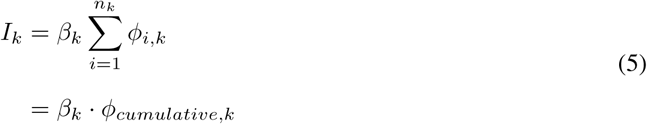

where *ϕ_cumulative,k_* = ∑*_i_ ϕ_i,k_* is the cumulative ZOI metric and is proportional to the ‘density’ of features in space (e.g. Panzacchi *et al*., 2015). The cumulative ZOI metric is easily calculated using geographical information systems, e.g. through neighborhood analysis, and can be rescaled to meaningful scales, such as the number of houses per km^2^. The same reasoning can be applied for variables represented as lines and polygons, such as roads, power lines, or mining sites; see the derivation of analogous equations in Appendix A.

## 3 Estimating the cumulative impact of multiple features

In the *cumulative ZOI approach*, the calculation of the potential ZOI (*ϕ*) is done before statistical analysis and *ϕ* based on different shapes and radii are considered as alternative predictor variables in a model of a biological response variable (Fig. 2). Among multiple ZOI predictors, only one or a few may be selected as an estimated ZOI through statistical fitting. Therefore, assessing the cumulative impact of multiple features and identifying the ZOI shape and size has been recast as a model selection rather than a parameterization problem (such as in Lee *et al*., 2020).

**Figure 2:**
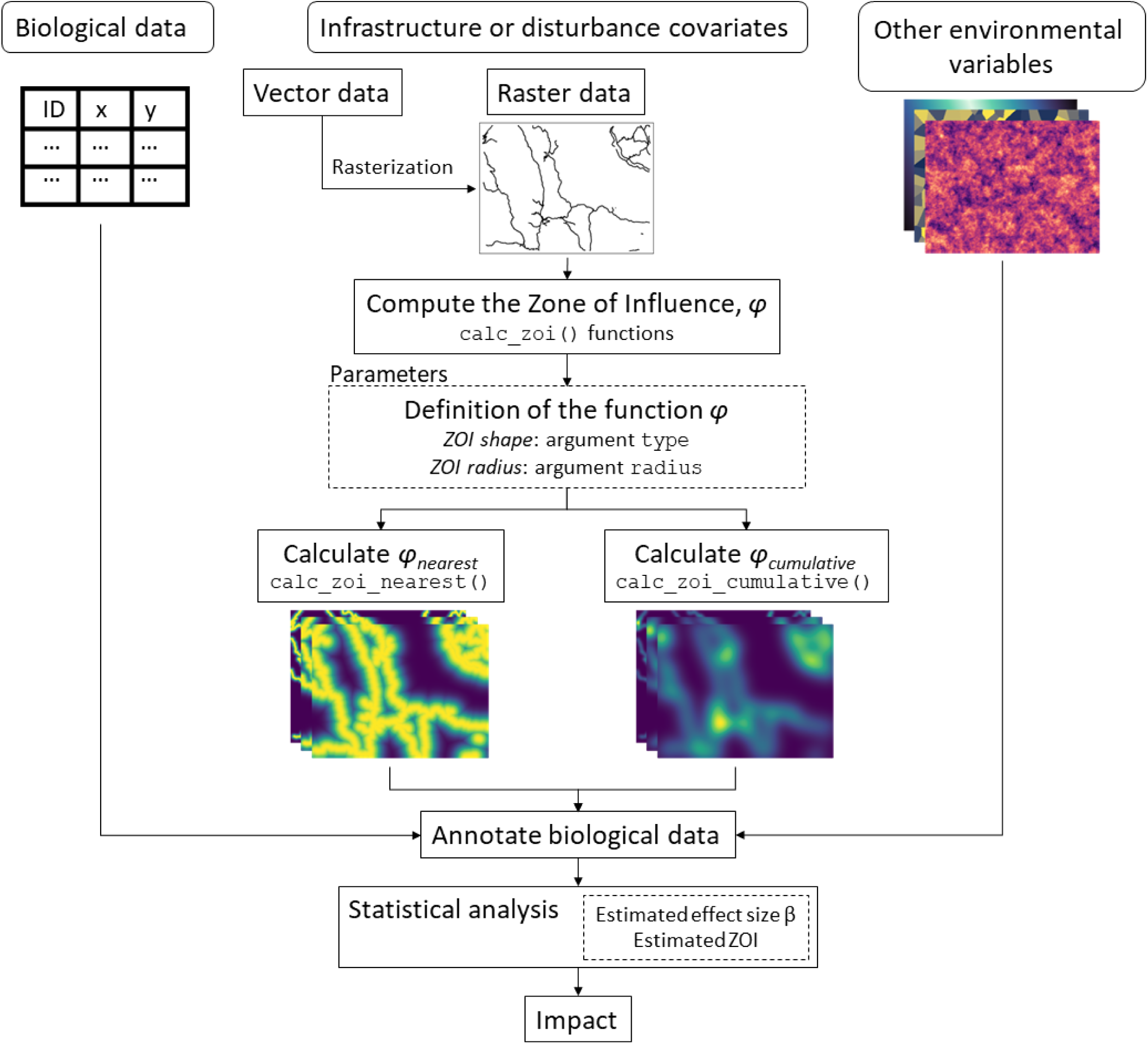
Workflow to calculate the ZOI *ϕ* and estimate the cumulative impact and ZOI radius of multiple infrastructure in the *cumulative ZOI approach*, using the `oneimpact` R package. The `calc zoi` functions use raster data describing infrastructure locations as input to calculate *ϕ_nearest_*and *ϕ_cumulative_* based on arguments describing the ZOI expected shapes and radii. Each output raster, which describes the ZOI defined by specific shapes and radii, is considered a different covariate. The output rasters, together with other environmental data, are then annotated to biological data and are analyzed to estimate *β* and *r* for each type of disturbance and calculate their total impact *I*.

The `oneimpact` R package has been developed to calculate *ϕ_nearest_* and *ϕ_cumulative_* through the `calc_zoi` functions that allow for ZOI defined by decaying functions of different shapes and radii and use raster maps representing infrastructure or other spatial variables as input (Fig. 2). Given spatially explicit biological data (e.g. species’ occurrence, abundance, or GPS positions of individuals), it is possible to spatially join the values of the ZOI (and, if relevant, of other environmental data) for all map pixels, thus producing a data set composed of biological response variables annotated with local and landscape-level predictors. The annotated data set is used to estimate the effect sizes *β* and the estimated ZOI and to evaluate the cumulative effects for different types of disturbances through statistical fitting of eq. 1 (Fig. 2). In the *cumulative ZOI approach*, the cumulative effect of multiple features is taken into account in the computation of the predictor ZOI variables. The approach is therefore applicable to a wide range of response variables and statistical modeling approaches. Therefore, the *cumulative ZOI approach* is useful for inferring cumulative impacts for a wide set of biotic or abiotic variables (similar to Lowe *et al*., 2022) related to different ecological processes (see Appendix B for examples). Similarly, when estimating the form and radius of *ϕ*, the approach can make use of more traditional model selection (Burnham & Anderson, 2002; Huais, 2018), penalized regression (Lee *et al*., 2020), or machine learning approaches (Pichler & Hartig, 2022), with different sampling designs and assumptions to account for spatial and temporal autocorrelation (see Northrup *et al*., 2022). A comprehensive review of such procedures is beyond the scope of this paper, but below we provide an example using model selection through AIC. Vignettes illustrating the `oneimpact` R package and the workflow in Fig. 2 are provided in https://ninanor.github.io/oneimpact/articles/.

## 4 When do *ϕ_nearest_* and *ϕ_cumulative_* represent similar spatial variation?

To correctly interpret *ϕ_nearest_*and *ϕ_cumulative_*it is important to understand under which conditions these two metrics are expected to represent similar gradients of spatial variation, and to be *de facto* equivalent. Similarities between the two variables depend on the spatial distribution of features as well as their ZOI, and might affect our ability to distinguish among their impacts. We simulated a set of landscapes (30 *×* 30 km^2^, 100m resolution) with a constant number of point features (*n* = 100) distributed according to different spatial patterns: regular, random, and clustered (Fig. 3; Appendix C). For each landscape we calculated *ϕ_nearest_* and *ϕ_cumulative_* assuming a range of ZOI radii (from 0.06% to 40% of the landscape extent), using a linear decay ZOI function (Fig. 1). We then compared the resulting spatial patterns of *ϕ_nearest_* and *ϕ_cumulative_* through Pearson correlation of the values of the two metrics at the same coordinates.

**Figure 3:**
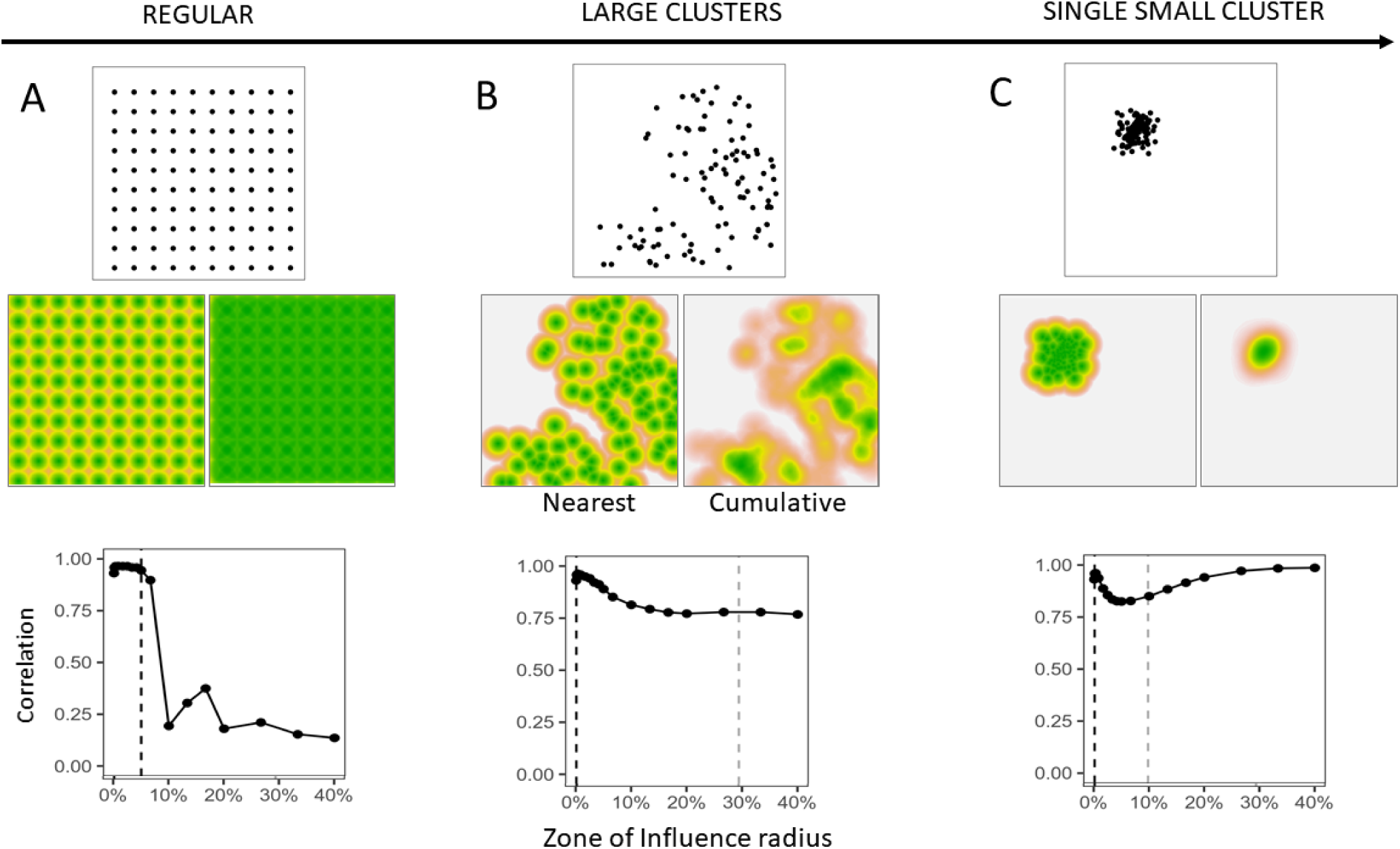
Representation of the ZOI of the nearest feature (*ϕ_nearest_*) and the cumulative ZOI (*ϕ_cumulative_*) in landscapes with point infrastructure spatially distributed in a gradient of clustering, from (A) a regular distribution to (B) a set of clusters to (C) only one cluster. The central panel illustrates *ϕ_nearest_* (left) and *ϕ_cumulative_* (right) when the ZOI radius *r* = 10% (3km) of the extent of the landscape. The lower panel shows the correlation between *ϕ_nearest_* and *ϕ_cumulative_* in each landscape, as the ZOI radius *r* increases. The dashed vertical lines show half the minimum distance between features (black), beyond which the ZOI of multiple features overlaps, and the size of the feature clusters (grey), beyond which the correlation stops decreasing.

When the minimum distance between features is greater than 2*r*, *ϕ_nearest_* and *ϕ_cumulative_* are identical (Fig. 3, black dashed vertical line; correlation = 1). This is because the ZOI of each feature is too restricted to overlap. When the ZOI radius increases, the effect of nearby features accumulate and the correlation between *ϕ_nearest_*and *ϕ_cumulative_*decreases (Fig. 3A,B, Figs C5 and C8). In addition, as the features get more aggregated (up to a limit with a single small cluster, Fig. 3C), the correlation between *ϕ_nearest_*and *ϕ_cumulative_* goes through a point of inflection as the ZOI expands, beyond which it increases with *r* (Figs C5D-F). The point where the correlation stops decreasing is related to the size of the clusters (grey dashed vertical line in Figs. 3B,C). For ZOI radii larger than the radius of the cluster, *ϕ_nearest_* and *ϕ_cumulative_* converge again and it may be hard to distinguish between the effect of each feature alone, regardless of the ZOI metric. At this point, the effect of a collection of features transforms into that of a ‘super-feature’ (e.g. urban areas instead of houses, wind parks instead of wind turbines).

## 5 Empirical demonstration: Impact of tourist infrastructure on reindeer habitat

### 5.1 Materials and methods

We evaluated the impact of tourist infrastructure on habitat selection of the Hardangervidda reindeer population in Norway during summer (Fig. 5). The Norwegian populations represent the last remaining wild mountain reindeer in Europe and are highly sensitive to human activities. In summer, their habitat is visited by tourists and hikers, and the area contains 14,154 private cabins, 26 large tourist cottages, and hundreds of kilometers of trails, in addition to other infrastructure (Fig. D2). We used GPS-tracking data from 115 female reindeer collected in the period of 1 July - 15 August 2001-2019 (see Panzacchi *et al*., 2015). Habitat selection was estimated using HSF in a use-availability setup, where each GPS location was compared with nine available random locations within the area (Fig. 5). All locations were annotated with environmental covariates (Fig. 2).

To account for bioclimatic and geographical variations we used the four first components from a principal component (PC) analysis (Bakkestuen *et al*., 2008). They correspond to gradients of (1) PC1 - continentality, (2) PC2 - altitude, (3) PC3 - terrain ruggedness, and (4) PC4 - solar radiation. We included a quadratic term for PC1 and PC2 to account for niche ‘optima’ (*sensu* Panzacchi *et al*., 2015). We also used a satellite-based land cover map with 25 vegetation classes, which were reclassified into 12 classes (Table D2). In this paper our aim was to illustrate the novel *cumulative ZOI approach*, and we thus needed to keep model fitting relatively simple, avoiding correlation between covariates. Therefore, we estimated the cumulative impact of only two of the main anthropogenic variables that occur in the core of the area inhabited by reindeer: private cabins and large tourist resorts. Note, however, that other infrastructure are present especially at the fringes of the study areas, and a proper assessment of the cumulative impact of all human activities on reindeer in Hargandervidda should account for all these anthropogenic variables (see Panzacchi et al., 2015).

For the raster layers describing private cabins and tourist resorts, we calculated both *ϕ_nearest_* and *ϕ_cumulative_* for a set of radii between 100m and 20,000m using the `calc_zoi()` functions of the `oneimpact` package (Fig. 2). For each infrastructure, we tested which of these four decaying functions best described the ZOI: threshold, linear decay, Gaussian decay, and exponential decay (Appendix A). To estimate reindeer habitat selection, we fitted HSFs (eq. 2) using binomial generalized linear models (Fieberg *et al*., 2021) with used and available locations as response and infrastructure, land cover, and bioclimatic variables as fixed effects. Model fitting consisted of two steps. We first fitted single-infrastructure models using a variable selection procedure (Burnham & Anderson, 2002) to find the most likely ZOI (shape, radius) for each infrastructure type. Single-infrastructure HSFs were fitted using the `multifit` function in R (Huais, 2018) and compared using AIC. Second, using the most likely ZOI from single-infrastructure models, we fitted multi-infrastructure HSF to assess the combined impacts of multiple types of infrastructure, as in Laforge *et al*. (2015). To quantify the impact of infrastructure, we applied eq. 3 using the *β* and *ϕ* estimated from the model with the lowest AIC. We then estimated habitat suitability by predicting the HSF (eq. 2) over the study area and rescaling the predicted values to the interval [0, 1]. For details on data, environmental covariates, modeling, and results, see Appendix D.

### 5.2 Results

Overall, single- and multi-infrastructure models including *ϕ_cumulative_* performed much better than models including *ϕ_nearest_* (Table D2). This presents strong evidence that the impact of private cabins and of tourist resorts accumulates over reindeer habitat, inducing the species to avoid these infrastructure types to a far larger degree when these infrastructures are clustered, as compared to when they are spaced far apart in the landscape. The most plausible model that included *ϕ_nearest_*was ranked in the 26^th^ position in the model selection results (Δ*AIC* = 921), and the most likely model that included the log-distance to the nearest feature was ranked 44^th^ (Δ*AIC* = 1197, Table D2). Interestingly, the most parsimonious model showed private cabins exerted a constant cumulative impact within a threshold ZOI of 10 km, while large tourist resorts followed an exponentially decaying cumulative ZOI with 20 km radius (Fig. 4; Table D2).

**Figure 4:**
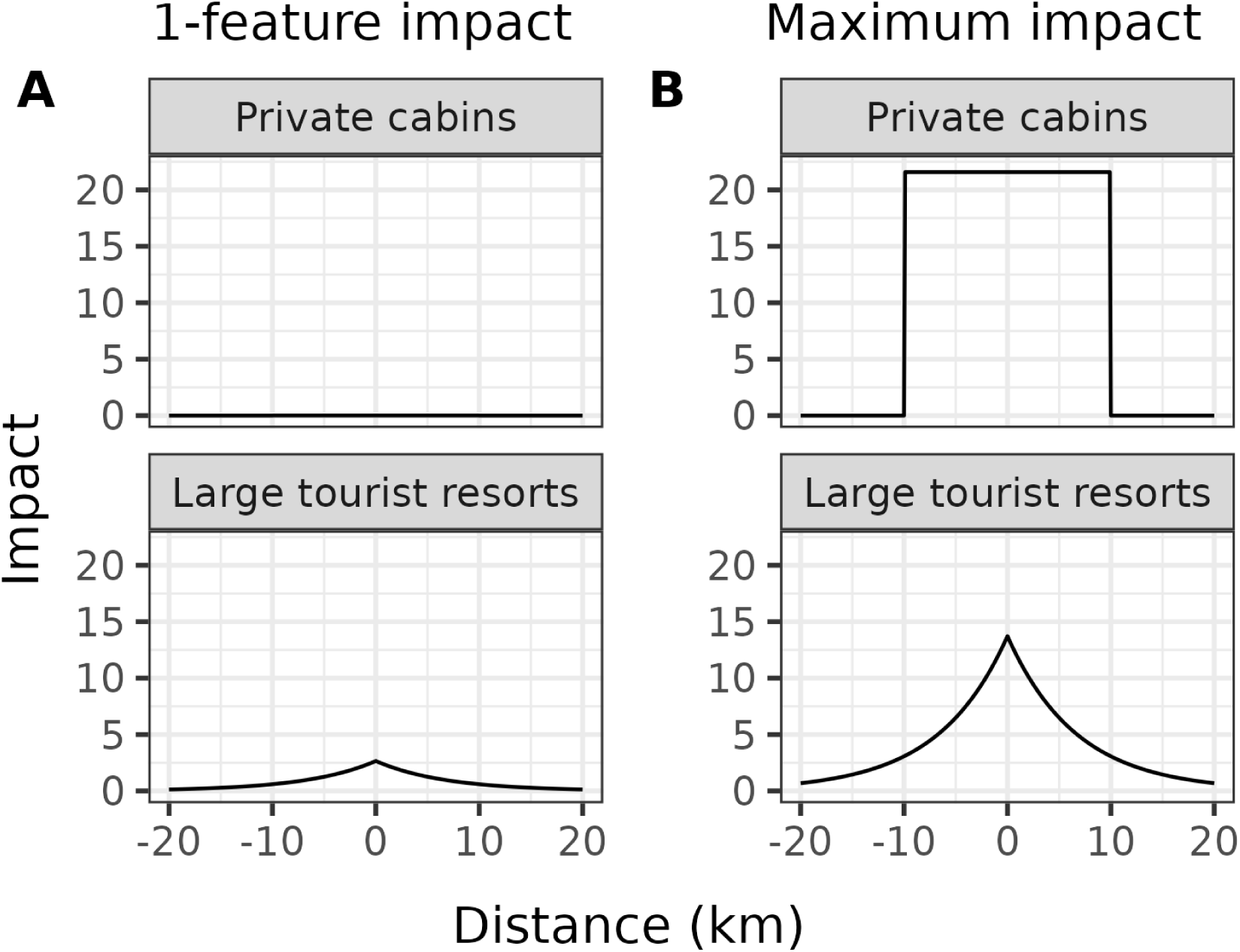
Impact of private cabins and public resorts considering (A) only 1 feature and (B) the maximum number of features of each type of infrastructure in the study area (2664 for cottages, 5 for cabins). The impact is the product of the effect size (*β*) and the cumulative zone of influence (*ϕ_cumulative_*). While the impact of one isolated private cabin is negligible (A), at their maximum densities the cumulative impact of “cabin villages” is higher than that of large tourist resorts (B).

The estimated effect size of a single private cabin (*β_cabin_* = *−*0.0081) was much smaller than that of a single tourist resort (*β_resort_*= *−*2.654; Fig. 4A, Table D3), as each private cabin is used by far fewer people (typically a family) compared to the tourist resorts (often used by hundreds of people). However, since private cabins can occur at higher densities in popular ‘cabin villages’, in some areas their impact is larger than that of a tourist resort. In the areas with the highest density of infrastructure in Hardangervidda – with 2664 private cabins and 5 tourist resorts – the impact of private cabin clusters is nearly twice that of tourist resorts (Figs 4B and 5). Following the interpretation of HSF coefficients from Fieberg *et al*. (2021), other conditions being constant, an addition of 330 private cabins (within 10 km) makes an area avoided by reindeer 14.4 times more strongly, what is nearly the same difference in avoidance a reindeer presents among two areas that differ in 1 tourist resort (within 20 km; Appendix D).

**Figure 5:**
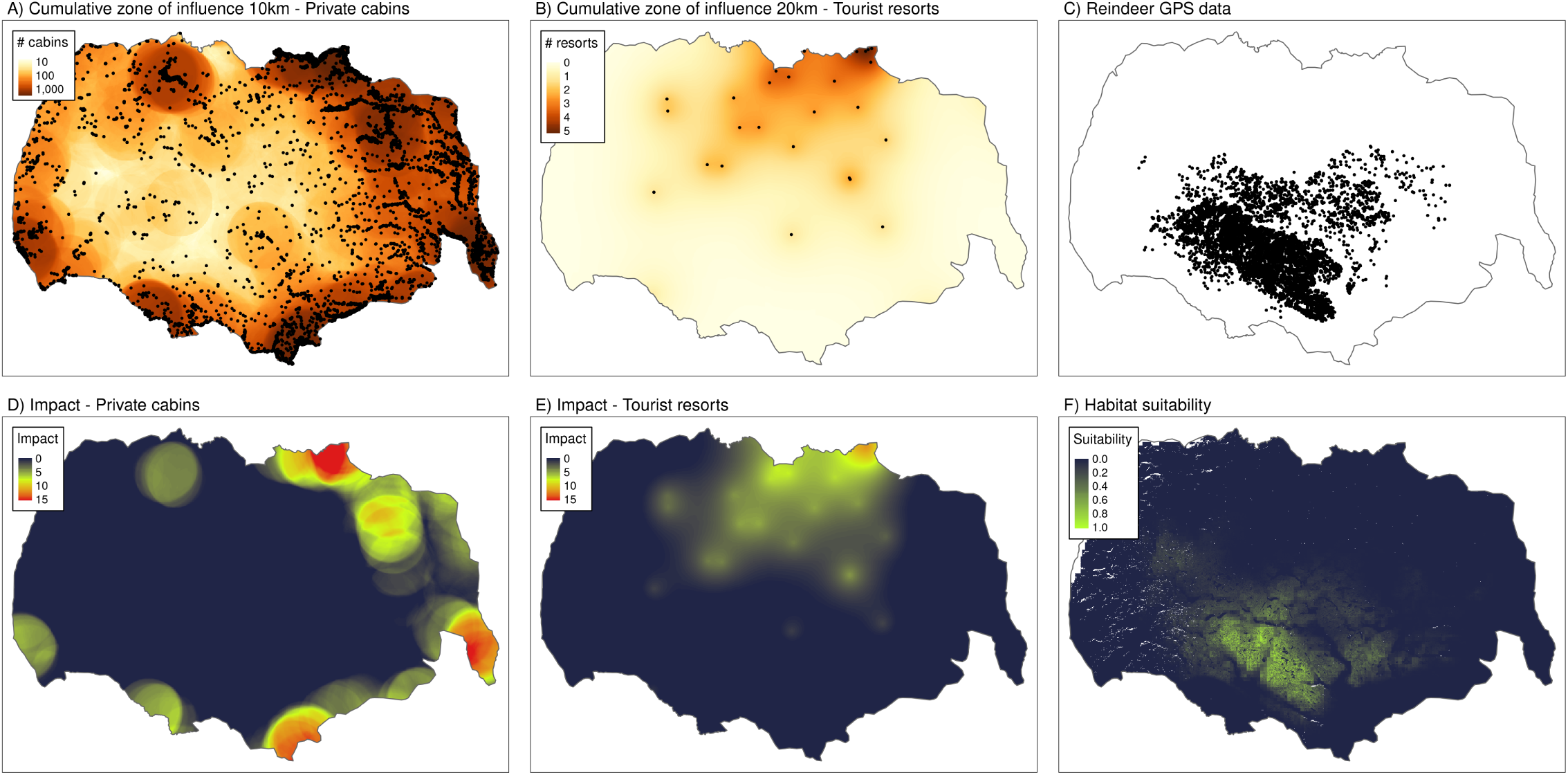
Maps illustrating the most parsimonious models for the estimated cumulative ZOI of private cabins (A, threshold model with 10km radius) and tourist resorts (B, exponential decay with 20km radius) and their estimated impacts (D, E) on reindeer habitat selection (F). These maps are shown alongside the GPS locations of reindeer during summer in the Hardangervidda wild reindeer area (C) and the predicted habitat suitability for reindeer, based on both the impact of cabins and cottages, as well as other environmental variables (F).

We predicted the cumulative impact of infrastructure in space by multiplying the effect size and *ϕ_cumulative_* (eq. 5). While the impact of private cabins rises 20 times in areas with the highest density of cabins, it does not increase more than 10 times for tourist resorts (Fig. 5). Due to the combined impact of infrastructure, and since reindeer avoided high densities of both infrastructure types at relatively large extents, areas of high habitat suitability for reindeer corresponded to those in which the cumulative impact of both infrastructure is low, which matches the locations used by reindeer, indicated through the GPS data (Fig. 5).

## 6 Discussion

There is an urge to correctly estimate and inform scientists, decision makers, and citizens about the past, current, and future impact of global land use changes on biodiversity (Laurance, 2018). Most decisions and regulations related to land use change are carried out with little statistical knowledge about cumulative impacts on ecosystems and species they affect (Johnson, 2011; Laurance & Arrea, 2017). Building upon previous frameworks (Johnson & St-Laurent, 2011) and concepts from landscape ecology literature to measure the ZOI of the nearest and of multiple features (*ϕ_nearest_*and *ϕ_cumulative_*), the *cumulative ZOI approach* provides tools to estimate such cumulative impacts on biodiversity.

Using simulations, we show that estimates based on nearest feature and cumulative impacts will be indistinguishable (i.e. *ϕ_nearest_* and *ϕ_cumulative_* are equivalent) in two extreme cases: (i) when features are distant in relation to the radius of their ZOI, they are too spaced for their impacts to accumulate; (ii) when features are highly clustered, they act as ‘super-features’ and the estimated ZOI will be large (e.g. building a new house increases the impact of an urban area little). In intermediate cases, which are common in real landscapes, cumulative impacts may be expected and should be accounted for. Although for illustrative purposes we focus here on the impact of two types of infrastructure on species’ habitat use, the approach is easily extended to cumulative impacts of and the full set of spatial features that affect a focal ecological process.

### 6.1 Applying the *cumulative ZOI approach* to assess impacts and habitat loss

Our empirical demonstration strongly supports the hypothesis of cumulative impacts of both private cabins and of tourist resorts on reindeer habitat use, with a ZOI of 10 and 20 km, respectively. These estimates fit with our knowledge of the distances typically covered by hikers relying upon their private cabins for day trips, and by tourists hikes from one tourist resort to another in multi-day trips. Our results show that while the impact of a single cabin is smaller than that of a tourist resort, the impact of several cabins, or cabin villages, can be far larger than that of a tourist resort (Figs 4, 5, D5). Similar results reporting a high impact of tourism and large ZOI for tourist facilities have been identified in a large number of studies with a variety of study designs over the last decades (Gundersen *et al*., 2019; Panzacchi *et al*., 2015, 2016, 2022; Polfus *et al*., 2011; Nellemann *et al*., 2001, 2010). It should be highlighted that the main aim of this study is to present the novel *cumulative ZOI approach*, and therefore we chose to minimize the landscape complexity and ignore a wide range of infrastructures known to impact reindeer habitat use, including trails, roads, railways, hydropower etc. The estimated impact of tourism presented here should therefore be considered as realistic, but indicative, as more precise estimates useful in applied contexts require considering also other anthropogenic features in the landscape (see e.g. Panzacchi *et al*., 2015, 2022). Specifically, accounting for other features correlated in space with private cabins and tourist resorts may reduce the ZOI estimated for these infrastructures.

Importantly, all our cumulative ZOI models (*ϕ_cumulative_*) had far more empirical support than models using nearest features (*ϕ_nearest_*). This suggests that models that consider the impact of the nearest feature only will often limit our understanding of the cumulative consequences of land use changes on biodiversity. The cumulative ZOI approach is a tool that helps analysts overcome this methodological limitation, and we believe will contribute to a more robust estimation of the impact of anthropogenic features and activities on ecological processes.

Studies measuring either distance or density of infrastructure are widespread in the literature, spanning movement ecology (Zeller *et al*., 2017), species distribution models (Panzacchi *et al*., 2015), population dynamics (Moraga *et al*., 2019), landscape genetics (Collevatti *et al*., 2020), species diversity and habitat models (Ficetola & Denöel, 2009), ecological interactions (Marjakangas *et al*., 2020), and assessment of abiotic conditions (Liu & Yang, 2018, Appendix B). However, in most cases, only one of the ZOI metrics was chosen to represent the effect of spatial features. Our approach allows flexible modelling of the ZOI for each type of feature and focal biological response variable. Furthermore, the method can and should be extended to estimate cumulative impacts of all anthropogenic features on focal species and ecological processes and is a crucial step towards comprehensive indicators of anthropogenic impacts.

The impact of anthropogenic features, such as infrastructure, is often assessed using study designs with some form of temporal or spatial replication. Examples of study designs with temporal replication are before-after or before-after-control designs that compare biological responses before and after the feature enters the landscape (Boulanger *et al*., 2021; Skarin *et al*., 2018; Dorber *et al*., 2023). Although these studies mimic experimental designs under controlled conditions, this assumption is rarely met in real landscapes, where many environmental variables may change simultaneously. In addition, the temporal replication of data both before and after tend to limited, allowing for little understanding of yearly variations and how anthropogenic features affect this variability. Other studies focus on spatial replications and study the response of species to different anthropogenic features across a wide range of environmental conditions.

In such studies, features may be isolated in some areas and aggregated in other areas, and landscape covariates are more likely to vary independently of the distribution of anthropogenic features. This design may help disentangle the effect of spatially correlated features and assess their average impact on ecological processes (e.g. Panzacchi *et al*., 2015), including on the species’ functional habitat networks (Van Moorter, in press). However, this spatial design is data demanding. Irrespective of the approach chosen to assess the impact of anthropogenic features, the *cumulative ZOI approach* offers a tool for understanding how the type and spatial configuration of features can lead to cumulative impacts and shape species’ spatial responses. Regardless of the study approach, investigating the occurrence of cumulative impacts is of paramount importance to avoid severe underestimations of habitat loss.

### 6.2 Assumptions, advantages, and limitations of the approach

The ZOI of anthropogenic features has been defined in different ways across studies, leading to different estimates and interpretations. Traditional studies used either a static, predefined buffer radius for each type of disturbance (Polfus *et al*., 2011), or post hoc analyses to define the ZOI after HSF fitting (Johnson *et al*., 2005; Plante *et al*., 2018). The *cumulative ZOI approach* proposed here consists of a model- and data-driven inference, which makes it robust and useful beyond the context of habitat use models, and particularly relevant in applied contexts where the ZOI differ for different ecological responses (Moraga *et al*., 2019). Importantly, it also allows the incorporation of uncertainty in the estimation of the ZOI, e.g. by computing confidence intervals through bootstrapping (Boulanger *et al*., 2021; Moraga *et al*., 2019).

It is worth noting that when functions of different shapes are chosen for modelling the ZOI, they imply different distribution of impacts inside the radius of the ZOI. In our example, the ZOI of private cabins indicated a homogeneous effect within an area with 10km radius (threshold function), while the impact of tourist resorts decayed exponentially within a 20 km ZOI radius. The latter implied that the impact decreases to half of its maximum value already 5 km from the feature (Fig. 4, Appendix A). Along the same lines, in a study of bird and insect abundances, Miguet et al. (2017) showed that the area affected by landscape variables can increase by a factor of 5 when using a distance-weighted influence measure (as used for tourist resorts in our example), in comparison to a threshold-based landscape measure (as used for private cabins in our example). We also note that the estimated values for *β* and ZOI radius can differ substantially between *ϕ_nearest_*and *ϕ_cumulative_*, depending on the spatial distribution of features. In our example, private cabins were abundant, the estimated ZOI radius for *ϕ_cumulative_* was 10 km and the effect size was small (Fig. D3), while the ZOI radius for *ϕ_nearest_* was only 1 km and the effect size was orders of magnitude higher (Fig. D3). However, such a difference in parameter estimates for *ϕ_cumulative_* and *ϕ_nearest_* was not observed for tourist resorts, which are scarce and sparsely distributed in the study area (Fig. D4).

In the `oneimpact` R package, the ZOI metrics are calculated before model fitting (Fig. 2). Here lies one of the main advantages of the approach – by precomputing the layers describing the cumulative impact of multiple features, we avoid the need for tedious iterative model fitting and complex estimation of parameters of nonlinear functions (Lee *et al*., 2020; Lowe *et al*., 2022). This facilitates the estimation of the ZOI for different feature types and eases model fitting for large datasets (e.g. Tucker *et al*., 2018) encompassing large study areas and fine-resolution spatial covariates. We believe that this makes the approach suitable for a wide range of ecological responses and study designs.

Our formulation of *ϕ* implies two main assumptions. First, for simplicity, the ZOI of each feature is assumed to be the same regardless of the density of points in an area. However, it may be more realistic to assume that the ZOI of a single or few features is smaller than that of a high-density cluster of features; for example, popular areas with clusters of tourist cabins are expected to be used by more people and potentially impact a wider area. Analogous calculations with variable radii have been implemented for decades in adaptive kernel density estimation (Worton, 1989), so our assumption can in principle be relaxed, and we encourage future developments in this direction to increase the local relevance of large-scale cumulative impact studies. Second, for simplicity, our formulation represents two extreme cases where either the nearest feature is the only one influencing the focal ecological process (*β_i_*= 0 for *i >* 1 in *ϕ_nearest_*), or all features affect the process equally (*β* is constant over all features in *ϕ_cumulative_*). More complex and realistic formulations could be derived extending eq. 3 to bridge the gap between the two extreme cases described above. Also, in principle, both *ϕ_nearest_*and *ϕ_cumulative_*can be included in the same statistical model, with the effect size estimated for each variable and a composite, cumulative impact inferred by combining the estimates.

## 7 Conclusions

The cumulative impact of multiple anthropogenic drivers is a major cause of the current unprecedented nature decline, with 75% of the land being significantly impacted, and 1 million species threatened with extinction (IPBES, 2019). However, comprehensive and robust scientific frameworks to study cumulative impacts are still under development, often leading cumulative impact assessments to the initiative of the analysts or impact assessors (Johnson, 2011). There is an urgent need to include more precise estimates of cumulative impacts in environmental impact assessments, sustainable land planning, and ecological studies. The *cumulative ZOI approach* takes this research field a step further and provides a framework to detect and estimate the magnitude and spatial extent of cumulative impacts of multiple spatial features in a wide range of ecological studies. The formulation presented here can be used to model cumulative impacts of anthropogenic features not only on species’ habitat selection, but on virtually all spatially explicit response variables, including population abundance (e.g. Beńıtez-López *et al*., 2010), species richness (e.g. Ficetola & Denöel, 2009), measures of biological diversity, community dynamics, and ecological processes such as movements. Therefore, the *cumulative ZOI approach* offers an opportunity to counter the widespread underestimation of the total impact of co-occurring anthropogenic disturbance factors in ecology, impact assessment, and land use planning.

## Acknowledgements

We thank C. Johnson for discussions on cumulative impact concepts, P.G. Blackwell for inputs on the equation notation, P. Dodonov, M.H. Vancine, and J. Nowosad for discussion around the implementation of ZOI metrics in R, F. Frassinelli for Github support, R. Muylaert, M. Grainger, the editor N. Cooper, and three anonymous reviewers for a critical review of the text, and E. Gurarie for the incentive and teachings on how to build an R package. We are grateful to several local projects allowing us to collect reindeer GPS data in Norway in the past decades and for NINA staff for managing them. The work presented here is the result of large research projects supported by the Research Council of Norway: OneImpact (n. 287925), RenewableReindeer (n. 243746), ProdChange (n. 255635), and GreenPlan (n. 326979). BBN, AS, PS, and MA were also supported by a grant from the Swedish Energy Agency within the research programme Vindval (grant n. 46780-1) and by project MineDeer (Vinnova project n. 2019-05191).

## Authors’ contributions

BBN, BVM, and MP conceived the idea and designed the methods, with contributions from TT, ASt, MA, and ASk; BBN, BVM, MP, TT, KL, and OS collected and provided the data; BBN, BVM, and MP analyzed the data and interpreted the results; BBN and BVM led the writing of the manuscript. All authors contributed critically to discussions and to the drafts, and gave final approval for publication.

## Data availability statement

GPS data is archived in Movebank (www.movebank.org) and can be accessed upon request. All environmental data was retrieved from public repositories. The `oneimpact` package is open and available at https://github.com/NINAnor/oneimpact, and all scripts used in the analyses are available in the Github repository https://github.com/bniebuhr/cumulative_zoi_paper.

## Supplementary Material

Appendix A. Defining and deriving the zone of influence for multiple infrastructure features

Appendix B. Applying the cumulative zone of influence approach to ecological studies

Appendix C. Comparing the zone of influence of the nearest feature with the cumulative zone of influence of multiple features

Appendix D. Cumulative impacts of infrastructure on reindeer space use: fitting habitat selection models

