## Appendix A for "Estimating the cumulative impact and zone of influence of anthropogenic features on biodiversity"

Defining and deriving the zone of influence for multiple features

**Abstract**

In this document, we provide additional details around the definition of the zone of influence (ZOI). First, we define the ZOI $\phi$ and present examples of ZOI functions with different shapes, parameterized with the ZOI radius. Second, we derive the ZOI of the nearest feature and the cumulative ZOI of multiple features of an infrastructure for types of infrastructure and spatial variables that are represented as lines (e.g. roads, power lines) or polygons (e.g. dams, mining sites). This complements the derivation for point-type features as presented in the main text of Niebuhr et al. *Estimating the cumulative impact and zone of influence of anthropogenic features on biodiversity.* Methods in Ecology and Evolution.

#### Introduction

In the main text, we defined the zone of influence (ZOI) and derived the estimation of the ZOI of multiple features of an infrastructure type under different assumptions, but only did so for anthropogenic features that can be represented as points, such as houses, tourist cabins, or wind turbines. Here we complement it by exemplifying ZOI functions with different shapes, parameterized on the ZOI radius $r$, and derive similar equations for linear infrastructure (e.g. roads, railways, trails, power lines) and infrastructure or landscape variables represented by polygons (e.g. mining sites, hydro dams, deforestation areas or polygons outlining specific land cover and land use types).

#### Definition of the Zone of Influence

As defined in the main text, the ZOI is the function $\phi$ that informs how the impact of a given feature decreases with distance. Formally, the ZOI $\phi = f(d, r)$ is any decay function that has a maximum value 1 where the feature is located, decreases towards zero as the Euclidean distance $d$ increases, and possibly vanishes at a given point, the ZOI radius $r$. Broadly speaking, the ZOI is characterized by its shape and radius. Here we use four functions with different shapes to evaluate how the impact decays with distance, even though in principle any function $\phi$ as defined above could be used. Below we write these four functions and show how the radius $r$ is defined in each one.

##### Functions with a well-defined ZOI radius

Some functions vanish for a certain non-infinite distance and therefore present well-defined radii. Here the ZOI radius $r$ represents the distance beyond which $\phi = 0$. We used two functions within this class: the threshold and the linear decay (Bartlett) functions.

###### Threshold function

The threshold function is defined as

$$\phi_{threshold}(d_{i,k}, r_k) = \begin{cases} 1 & \text{if } d_{i,k} < r_k \\ 0 & \text{if } d_{i,k} \geq r_k \end{cases}$$

where $d_{i,k}$ is the distance to the feature $i$ of type $k$ and $r_k$ is the radius of the ZOI for the features of type $k$. Then, $\phi_{threshold}$ represents a constant impact of each feature until $d = r$ and is similar to what has generally been used when buffer zones are created around anthropogenic features to calculate derived measures.

Fig. A1 shows a plot of $\phi_{threshold}$ with $r = 10$ km.

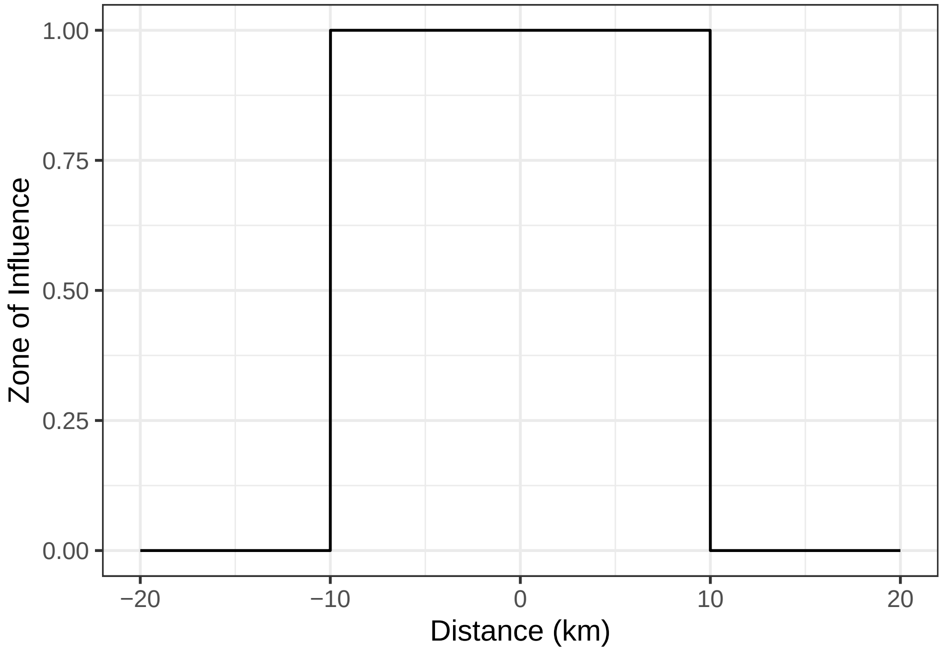


Figure A1: Illustration of a threshold ZOI function with radius = 10 km, for a feature located at the origin.

**Linear decay function**

The linear decay function, also called tent decay function or “Bartlett” filter (Harris, 1978), corresponds to 1 at the feature location and decreases linearly with the distance from it, until it reaches zero at a given distance, defined here as the ZOI radius $r$. Formally, the linear ZOI function $\phi_{linear}$ is defined as:

$$\phi_{linear}(d_{i,k}, r_k) = \begin{cases} 1 - \frac{d_{i,k}}{r_k} & \text{if } d_{i,k} < r_k \\ 0 & \text{if } d_{i,k} \geq r_k \end{cases}$$

where $d_{i,k}$ is the distance to the feature $i$ of type $k$ and $r_k$ is the radius of the ZOI for features of type $k$.

Fig. A2 shows a plot of $\phi_{linear}$ with $r = 10$ km.

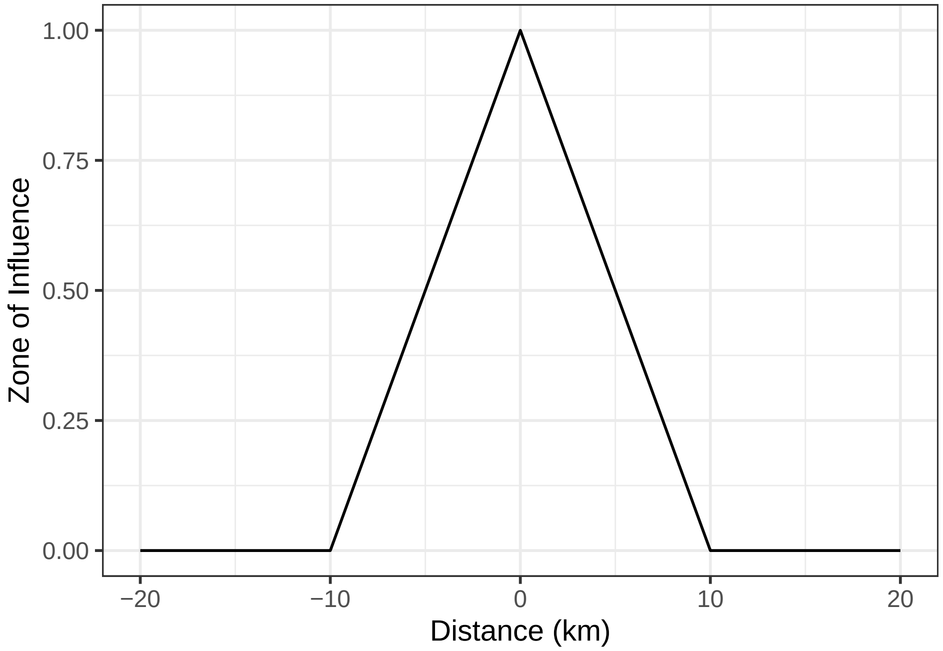


Figure A2: Illustration of a linear decaying ZOI function with radius = 10 km, for a feature located at the origin.

#### Functions that do not vanish with distance

Some functions decrease but do not vanish as the distance from anthropogenic features increase. In these cases we define the ZOI radius $r$ as the distance at which the ZOI decreases to $\phi = \phi_{limit}$, an arbitrary small ZOI value beyond which the influence of the feature is considered to be negligible. In these cases, the ZOI definition needs an extra parameter and is defined as $\phi = f(d, r, \phi_{limit})$. We used two functions within this class: the exponential decay and the Gaussian decay functions. For both we set $\phi_{limit} = 0.05$ for all features.

##### Exponential decay function

The exponential decay function is defined as

$$\phi_{exp}(d_{i,k}, r_k, \phi_{limit_k}) = \exp(-\lambda_{r_k} d_{i,k})$$

where $d_{i,k}$ is the distance to the feature $i$ of type $k$ and the decay parameter $\lambda_{r_k}$ is defined in terms of $r_k$ and $\phi_{limit_k}$:

$$\lambda_{r_k} = \frac{\ln(1/\phi_{limit_k})}{r_k}$$

Here ln is the logarithm with base $e$. Fig. A3 shows a plot of $\phi_{exp}$ with $r = 10$ km.

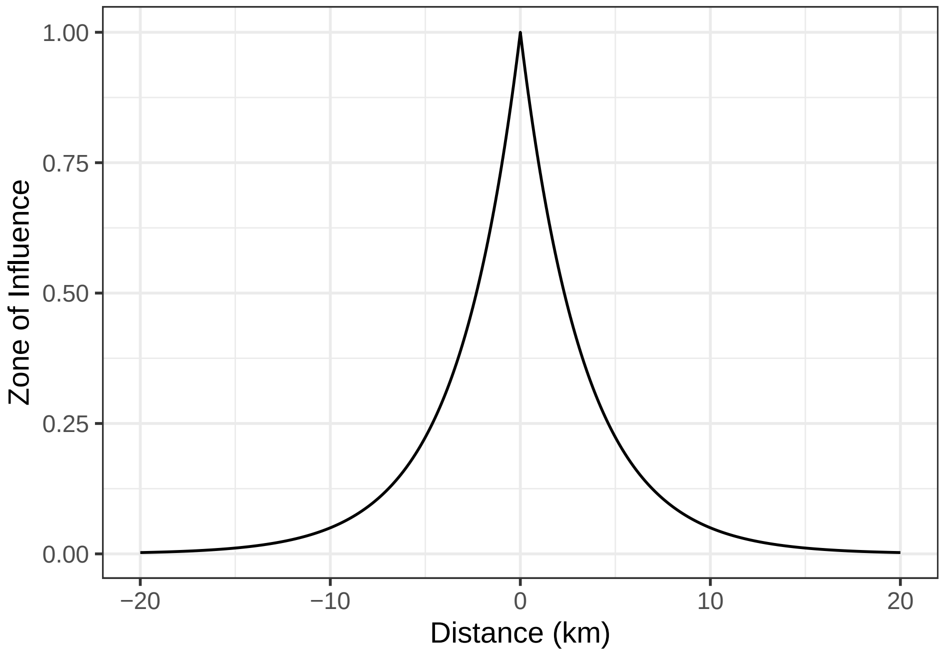


Figure A3: Illustration of an exponential decaying ZOI function with radius = 10 km, for a feature located at the origin.

**Gaussian decay function**

The Gaussian (or half-normal) decay function is defined as

$$\phi_{Gauss}(d_{i,k}, r_k, \phi_{limit_k}) = \exp(-\lambda_{r_k} d_{i,k}^2)$$

where $d_{i,k}$ is the distance to the feature $i$ of type $k$ and the decay parameter $\lambda_{r_k}$ is defined in terms of $r_k$ and $\phi_{limit_k}$:

$$\lambda_{r_k} = \frac{\ln(1/\phi_{limit_k})}{r_k^2}$$

In Fig. A4 we visualize $\phi_{Gauss}$ with $r = 10$ km.

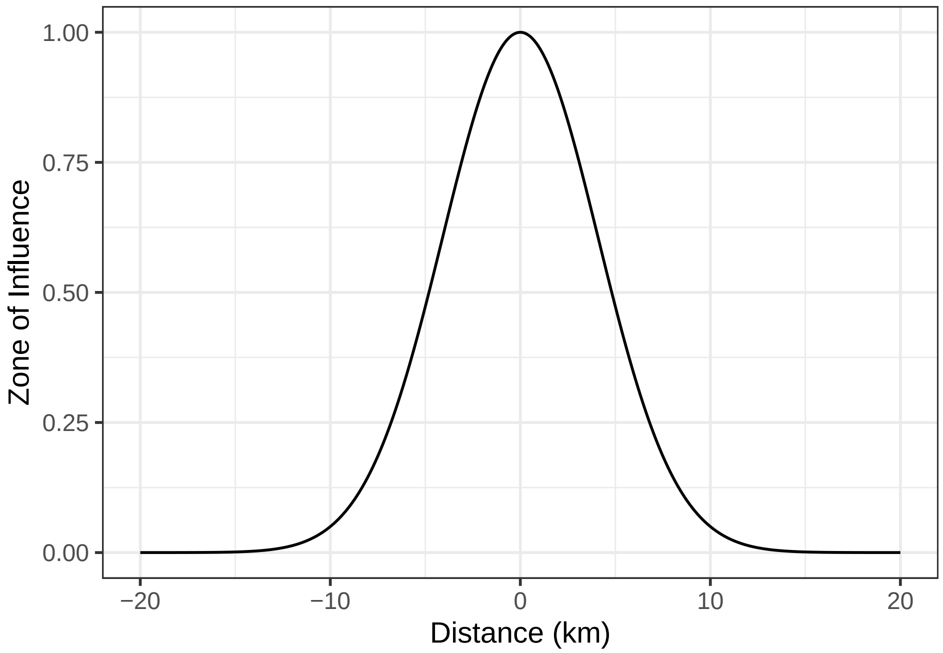


Figure A4: Illustration of a Gaussian decaying ZOI function with radius = 10 km, for a feature located at the origin.

Notice that, even though the ZOI radius $r$ is defined for all the functions, the change in their shape modifies the interpretation of how the ZOI changes with distance (see also Fig. 1 in the main text).

### Derivation of zones of influence for multiple anthropogenic features of a given type

We start from the same baseline presented in the main text: we use as an example of statistical model a habitat-selection function (HSF; Fieberg et al., 2021). Copying here eq. 2 from the main text, the HSF is defined as:

$$w(\mathbf{X}) = \exp\left(\beta_0 + \overbrace{\beta_1 X_1}^{\text{A) Infrastructure type 1}} + \overbrace{\beta_2 X_2}^{\text{B) Infrastructure type 2}} + \underbrace{\beta_{12} X_1 X_2}_{\text{D) Interaction infrastructure types 1 and 2}} + ... + \overbrace{\beta_k X_k}^{\text{C) Infrastructure type k}}\right) \quad (1)$$

where $\mathbf{X} = X_1, X_2, ..., X_k$ correspond to $k$ different types of infrastructure or other anthropogenic features and $\beta_k$ represents the effect size of the impact of spatial features of type $k$.

#### ZOI metrics

As shown in the main text, when multiple features of the same type are present in the landscape, we can calculate two ZOI metrics: the ZOI of the nearest feature only ($\phi_{nearest}$) and the cumulative ZOI of multiple features ($\phi_{cumulative}$).

We assume that in the landscape there are $n_k$ features of the same type $k$, and let the influence of the feature $i$ of a feature $k$ be a ZOI function $\phi$ as defined above. As we show in the main text, for point-like infrastructure we can sum the effect of each feature, so that the impact terms in equation 1 become:

$$I_k = \beta_k X_k = \sum_{i=1}^{n_k} \beta_{i,k} \phi_{i_k}. \tag{2}$$

Likewise, for linear infrastructure we can integrate the term

$$I_k = \beta_k X_k = \int_L \beta_k(l)\ \phi_k(l; d_{i,k}, r_k) dl \tag{3}$$

where $dl$ denotes the length on an infinitesimal infrastructure segment and $L$ is the total length of linear infrastructure of type $k$ in the study area. For polygon variables we can similarly integrate the term, but on two dimensions $(x, y)$:

$$I_k = \beta_k X_k = \iint_\Omega \beta_k(x, y)\ \phi_k(x, y; d_{i,k}, r_k)\ dx\ dy \tag{4}$$

where $\Omega$ is the whole study area.

#### Zone of influence of the nearest feature

To estimate the ZOI of the nearest feature alone, we showed that, for point-like features, one generally considers $\beta_i = 0$ for all $i > 1$ (where the features are ordered by increasing distance), i.e., only the nearest feature is assumed to influence a given site. We can make equivalent assumptions for linear and area infrastructure by considering only the nearest (orthogonal) segment of linear infrastructure or the closest edge of the closest polygon to influence processes in a given location. In such a way, their influence is similar to that of a point. As a consequence, eq. 4 from the main text keeps valid to represent the influence of the nearest feature for linear and polygon variables:

$$\begin{aligned} I_k &= \beta_{1,k} \cdot \phi_{1,k} \\ &= \beta_k \cdot \phi_{nearest_k}. \end{aligned} \tag{5}$$

Here $\phi_{nearest}$ can be set to represent the distance to the nearest feature (or some transformation of that, such as the log-distance to the nearest feature), as it is most common in the literature. However, our formulation with the definition of the set of functions $\phi$ to represent the ZOI allow for more a general representation of the impact of the nearest feature, while still allowing the estimation of the shape and radius of the zone of influence.

#### Cumulative zone of influence

To derive the cumulative ZOI of multiple features of a given type, for point-like features we assumed that all features exerted the same influence, so that $\beta_{1,k} = \beta_{2,k} = ... = \beta_k = \beta$. If we do similar assumptions for lines and polygons – that all infinitesimal segments $dl$ present the same influence over $L$ and that all pieces of area of the polygons present similar influence around them, eq. 5 from the main text remains valid for these types of variables:

$$\begin{aligned} I_k &= \beta_k \int_L \phi_k(l)\ dl \\ &= \beta_k\ \phi_{cumulative_k} \\ &\text{and} \\ I_k &= \beta_k \iint_\Omega \phi_k(x, y)\ dx\ dy \\ &= \beta_k\ \phi_{cumulative_k}. \end{aligned} \tag{6}$$

The only difference here regards how the cumulative ZOI function $\phi_{cumulative}$ is defined – as a summation for point features (eq. 5 in the main text) or as an integral over 1 or 2 dimensions for linear and area features (eq. 6).

### Computation of the cumulative zone of influence

In theory, the calculation of the cumulative zone of influence function $\phi_{cumulative}$ for a given study area $\Omega$ requires the computation of eq. 4 from the main text or eq. 6 for each infrastructure feature, which might be difficult and computationally challenging for large areas with many anthropogenic features spread non-regularly in space. To overcome that limitation, we implemented this calculation by constructing filters (or weighing matrices; see Miguet et al., 2017) that follow an assumed decaying function (such as the ZOI functions described above) and used them on neighborhood analyses over the input infrastructure layers (see the vignettes of the `oneimpact` R package for an illustration of how this approach is applied). This is analogous to the approaches presented by Miguet et al. (2017) and Gilleland (2013). Even though we do not demonstrate it here, for non-vanishing function (e.g. exponential or Gaussian decay) when the size of the filter is set properly to include most the area within which $\phi > 0$ (i.e. $\phi_{limit}$ is small enough), the result of applying eq. 4 of the main text or eq. 6 presents a correlation $> 0.99$ with the result of the neighborhood analysis.
