## Appendix B for "Estimating the cumulative impact and zone of influence of anthropogenic features on biodiversity"

#### Applying the cumulative zone of influence approach to ecological studies

**Abstract**

In this document, we provide examples of potential applications of the cumulative zone of influence approach in ecological studies with different contexts, biological measurements, and studied taxa. This is a supplementary information for Niebuhr et al. *Estimating the cumulative impact and zone of influence of anthropogenic features on biodiversity.* Methods in Ecology and Evolution.

### Introduction

In the main text, we present an application of the cumulative Zone of Influence (ZOI) approach on habitat selection, using reindeer as a model study. As mentioned there, the approach might be applied way beyond animal movement ecology studies. In principle, the approach is useful in any context in which predictor variables (sources of disturbance, infrastructure, or any other spatial variable) vary in space and are expected to affect the biological response variable or process being studied beyond their actual limits. This includes but is not limited to those studies in which either the distance to the nearest feature is computed and used as a predictor (e.g. distance to roads in Torres et al. 2016, or distance to forest edges in Martello et al. 2022) or those in which the density or amount of a landscape variable is computed (e.g. Jackson & Fahrig 2015, Miguet 2016, Martin 2018). Review literature shows that the so-called multiscale approaches have been applied for a different set of taxa (MacGarigal et al. 2016, Martin 2018) using distinct biological measures for a wide range of ecological processes (Martin 2016). Here we present examples of contexts in which our approach might be useful to estimate cumulative impacts and zones of influence, and provide studies to illustrate questions in which these dimensions could have been evaluated.

### Illustrative examples

In table B1, we present some examples of how the cumulative impacts of landscape disturbance and human infrastructure could be inferred on different ecological processes, using distinct types of data, for different response variables and organisms. For instance, studies on movement and behavioral ecology may evaluate how movement rates (e.g. Hansbauer et al. 2008) or the probability of an animal being in a given behavioral state (e.g. Morales et al. 2004) depend on the influence of disturbances in the landscape, and this influence might be measure through either the ZOI of the nearest feature or the cumulative ZOI metrics. Such response measures can based on different types of data (e.g. locations recorded through GPS, VHF, or other devices, capture-mark-recapture and camera trap data) for different organisms, from animals who move actively (e.g. Ramos et al. 2020) to seeds, fungal spores, or other propagules that are displaced by animals, wind, or other agents (e.g. Carlo et al 2015, Norros et al. 2012).

Likewise, in the context of populations and communities, for example, the ZOI metrics related to disturbance sources might be computed around sampling points, considering multiple radii, and used to fit models estimating their effects on population-level metrics (growth rate, survival/mortality, fecundity; e.g. Cerqueira et al. 2021) for basically any species or on community-level metrics (species diversity metrics, species abundance, co-occurrence probability, interaction network metrics; e.g. Torres et al. 2016).

More examples might be found in Table B1. We also provide in the table a few references illustrating studies in which a similar approach was used or could be adapted to incorporate an evaluation of cumulative impacts. In most of these studies, though, only one of the metrics was used - the distance to the nearest infrastructure or land cover type, related to the ZOI of the nearest feature, or the density or amount of landscape variables, related to the cumulative ZOI as presented here.

Looking more broadly, the approach can still be useful in other contexts in which landscape variables measures at different scales might play a role, e.g. in environmental research involving habitat structural and abiotic measures (Lowe et al. 2022). For instance, the amount of different pollutants in landscapes with different amounts of forest, urban area, and roads could be evaluated, or the amount of pesticides identified in the water could be tested against the proximity and cumulative influence of agricutural and forested areas to the sampling sites (e.g Liu and Yang 2018).

Table B1: Examples of cases in which the cumulative impact approach could be applied, considering different ecological processes, response variables, types of data, statistical models, and organisms studied. The last column shows some references in which a similar approach was used or could have been applied.

| Ecological process | Response variable | Type of data | Statistical model | Taxon | Reference |
|---|---|---|---|---|---|
| Movement ecology | Movement rate, step length, dispersal distance, turning angles | GPS, VHF, Capture-mark-recapture | Generalized linear (mixed) models [glm/glmm] | Animals, seeds, fungi, other propagules | [7, 8, 10, 11, 13] |
| Habitat selection Resource- or step- selection functions | Presence, given availability | GPS | Binomial glm/glmm, conditional regression | Animals | [6, 14, 15, 16, 17] |
| Ecological niche/ Species distribution models | Presence only, presence-background | Ocurrence, camera-trap | glm, MaxEnt, RandomForest | Any organism | [16, 18, 19, 20, 21] |
| Population dynamics | Population size, growth rate, survival, mortality, fecundity | Occurrence, count, distance-sampling, camera-trap | glm/glmm, occupancy models, survival models | Any organism | [5, 12, 22] |
| Community dynamics | Species (co)occurrence, species occurrence probability, species richness, species abundances, diversity indices | Multi-species occurrences or abundance | Joint species distribution models | Multiple | [1, 2, 23, 24] |
| Ecological interactions | Interaction network metrics, probability of interaction | Focal sampling of interactions | glm/glmm | Any type of interaction (e.g. throphic, mutualism, comensalism) | [25, 26, 27, 28] |
| Genetics | Kinship, effective population size, genetic diversity, inbreeding coefficients, genetic differentiation between populations | Genetic samples from multiple populations or areas | Multiple | Any organism | [16, 29] |
| Structural habitat, fragmentation, pollution | Habitat edge influence measures, amount of pollution particles, bioaccumulation, concentration of pesticides in the water | Multiple | Multiple | NA | [30, 31] |

### Using the approach

To apply the cumulative ZOI approach in these different contexts, we might start by following the same workflow presented in Fig. 2 of the main text. First, using the spatial variables of interest, we should compute the two ZOI metrics for multiple radii and shapes, as relevant for the empirical case. Once this is done, they can be used to annotate the biological data considering their spatial locations or sampling sites. Given the table with response and predictor variable data is ready, equation 1 from the main text might be adapted to look for the relationship between the sampled biological response variables and the ZOI and other spatial or non-spatial variables. Here the analyst might use whatever modeling approach suits the data and context better; the possibility of cumulative effects of each single spatial variable type and their scale of effect or zone of influence are already considered since they were used to compute the ZOI predictor variables. This means that, in this formulation, it is not necessary to change the model specification to assess the cumulative impacts of multiple features of a given type of variable. More details on how this is done in practice are shown in the vignettes of the `oneimpact` R package.
