## Appendix C for "Estimating the cumulative impact and zone of influence of anthropogenic features on biodiversity"

Comparing the zone of influence of the nearest feature with the cumulative zone of influence of multiple features

**Abstract**

In this document, we aim to illustrate in which conditions the zone of influence (ZOI) of the nearest feature alone ($\phi_{nearest}$) and the cumulative zone of influence of multiple features ($\phi_{cumulative}$) diverge or converge. First we explore it by simulating landscapes with different spatial configurations of anthropogenic features. Second, we calculate these metrics and compare them for tourist-related infrastructure in the Hardangervidda wild reindeer area in Southern Norway. Understanding how these ZOI measures change across landscapes is important to aid the interpretation of their effects. This document complements the arguments presented in Niebuhr et al. *Estimating the cumulative impact and zone of influence of anthropogenic features on biodiversity.* Methods in Ecology and Evolution.

**Additional remarks** **10**

### Introduction

There are several dimensions that underlie the assessment of the impacts of anthropogenic infrastructure on wildlife. When performing environmental impact assessments, one aims not only to to find which factors affect wildlife and how strongly, but also (i) at which spatial scale there are impacts (the Zone of Influence, ZOI) and (ii) how these impacts sum and interact when (as it is often the case) there are multiple infrastructure and vectors of landscape modification. In the main text of this manuscript, we argue that the cumulative ZOI ($\phi_{cumulative}$) – proportional to the spatial density of infrastructure features – might better represent the spatial variation of the cumulative impact of multiple features in the landscape than considering the ZOI based on the nearest feature only ($\phi_{nearest}$). How similar the two ZOI metrics are and whether one can distinguish between these them, however, might vary depending on the number of features in the landscape, how these features are distributed in space, and what is the ZOI of each of those features.

In this document we aim to assess in which conditions $\phi_{nearest}$ and $\phi_{cumulative}$ converge and diverge, to allow a better interpretation of their effects on biological response variables in ecological contexts. We first simulate landscapes with point-type infrastructure spread following different patterns, calculate both ZOI metrics, considering multiple ZOI radii, and assess when and how these variables might represent different sources of spatial variation. Second, we do the same for a real landscape, the Hardangervidda wild reindeer area in Southern Norway, which is the stage for the the habitat selection analyses presented in the main text and in Appendix D.

### Zone of influence metrics in simulated landscapes

#### Simulating landscapes

We start by simulating landscapes using point-type infrastructure. They could represent the spatial location of houses, cabins, or wind turbines, for example. We set 30x30 $km^2$ landscapes and simulate points following different spatial patterns: regular, random, and clumped distribution of points. For the clumped distribution, we set four landscapes by varying the number of clusters and the size of the clusters: point features are spread on either 5 or only 1 focal cluster, and we choose two possible radii of these clusters, corresponding to one more and one less clumped point distribution (radius = 5% [1.5 km] and 15% [4.5 km] of the extent of the landscape[1]). In such a way, we aim to represent a gradient of spatial clustering of the simulated point features, to evaluate how the ZOI metrics vary within this gradient. To keep the interpretation simple, we fix the total number of features in the landscape to $n_k = 100$. Yet, we also performed simulations with different numbers of features ($n_k = 10$) and the results are qualitatively similar.

The landscapes were simulated using the function `set_points()` from the `oneimpact` package. Fig. C1 shows six example landscapes simulated in this gradient of clustering.

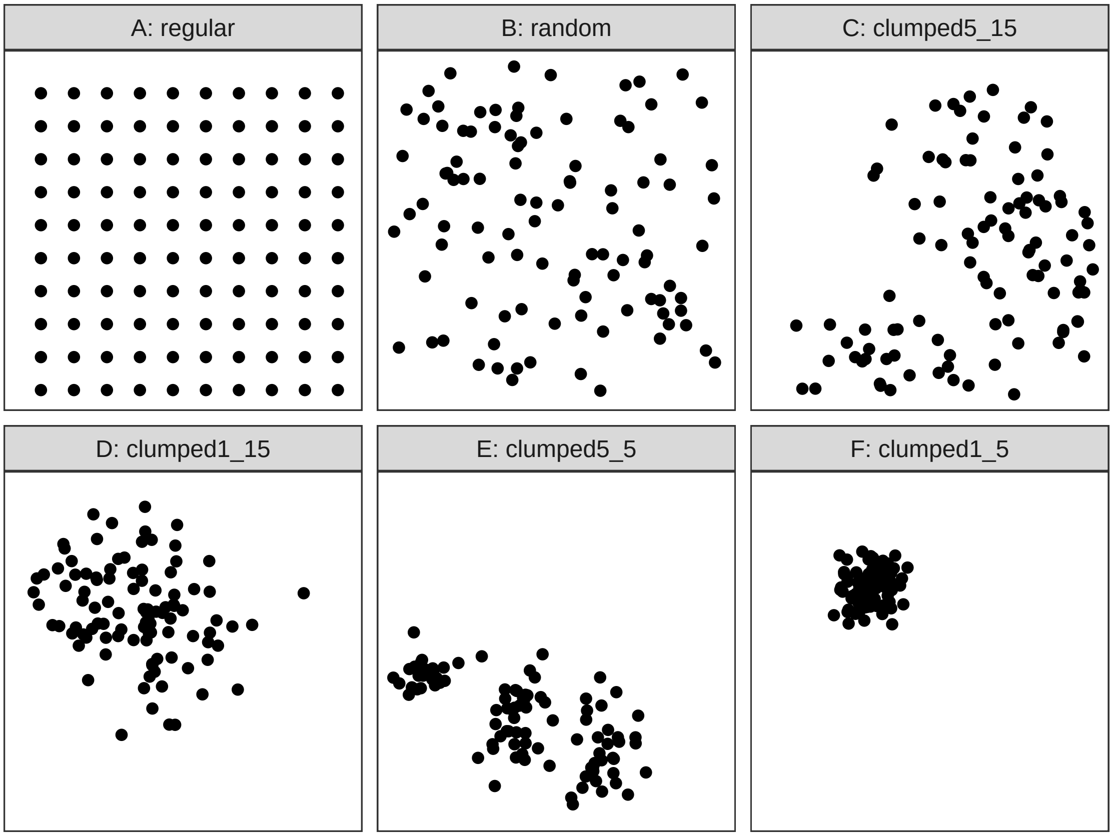

Figure C1: Six example simulated landscapes with points distributed in a gradient of clustering, from regular to random to clustered. The names of the landscapes with clustered distributions (C-F) also present the number of clusters and the radius of each cluster. For instance, "cluster1_15" represents the clumped distribution with 1 single cluster with radius = 15% of the landscape extent.

#### Calculating the zone of influence metrics

For each of the simulated landscapes, we calculate both $\phi_{nearest}$ and $\phi_{cumulative}$ with ZOI radii varying from ~ 0.06% to 40% of the landscape extent (i.e., from 20 m to 12 km, given our landscapes are squares of 30 km extent).

[1] To be precise, these values correspond to the standard deviation of the distance from points to the center of the cluster, simulated using a bivariate Gaussian distribution. To ease the interpretation, however, we call them radii of the clusters.

For the comparison presented here, we computed both ZOI metrics using a linear decay ZOI function (see Fig. 1 in the main text), for which the ZOI can be defined only based on the ZOI radius $r$ (see Appendix A). The ZOI rasters were computed using the function `calc_zoi()` from the `oneimpact` package in R.

#### Visualizing the zone of influence metrics

To visualize qualitatively how different are $\phi_{nearest}$ and $\phi_{cumulative}$, we plot them for different spatial configuration of features, considering different ZOI radii. We see in Figures C2, C3, and C4 the two metrics for $r = 1.6\%$, 5%, and 10% of the landscape extent (500, 1500, and 3000 m). When the ZOI radius $r$ is small compared to the landscape extent, the two ZOI metrics are similar and, in the computation of $\phi_{cumulative}$, there is interaction between the ZOI of multiple features only for very clustered feature distributions (Fig. C2). As $r$ increases, however, going beyond the minimum distance between features, the ZOI from different features add up in the calculation of $\phi_{cumulative}$ and the two ZOI metrics diverge (Fig. C3 and C4). Notice that the values in the color scale change between ZOI metrics for the different conditions of feature spatial configuration in Figs. C2-C4.

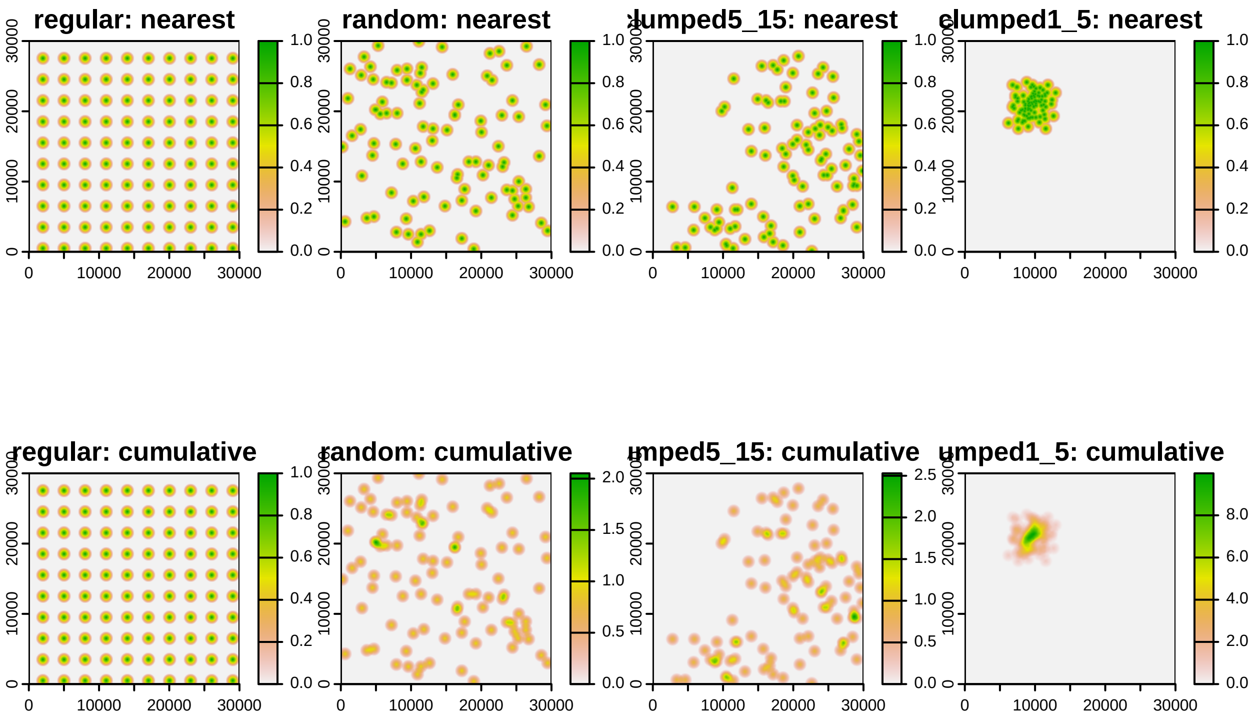

Figure C2: Illustration of the zone of influence of the nearest feature and the cumulative zone of influence of features for 4 landscapes in a gradient of clustering of the point infrastructure, for ZOI radius = 1.66% of the landscape extent (500 m).

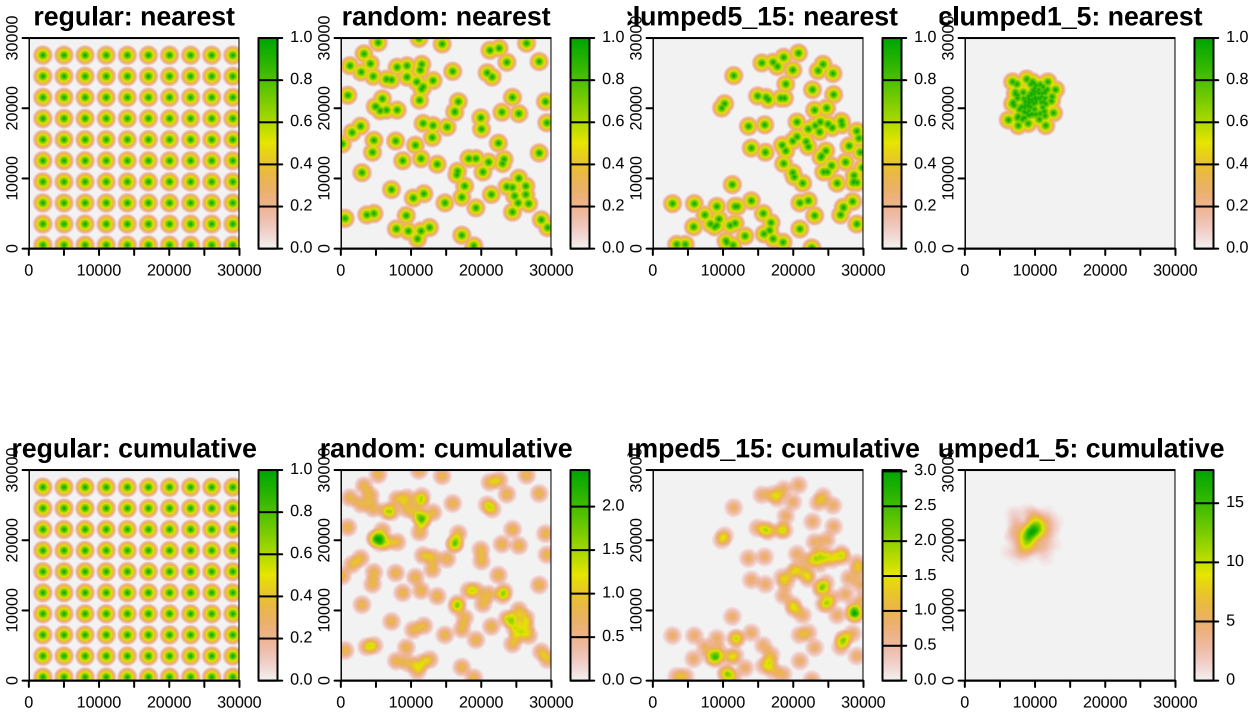

Figure C3: Illustration of the zone of influence of the nearest feature and the cumulative zone of influence of features for 4 landscapes in a gradient of clustering of the point infrastructure, for ZOI radius = 5% of the landscape extent (1500 m).

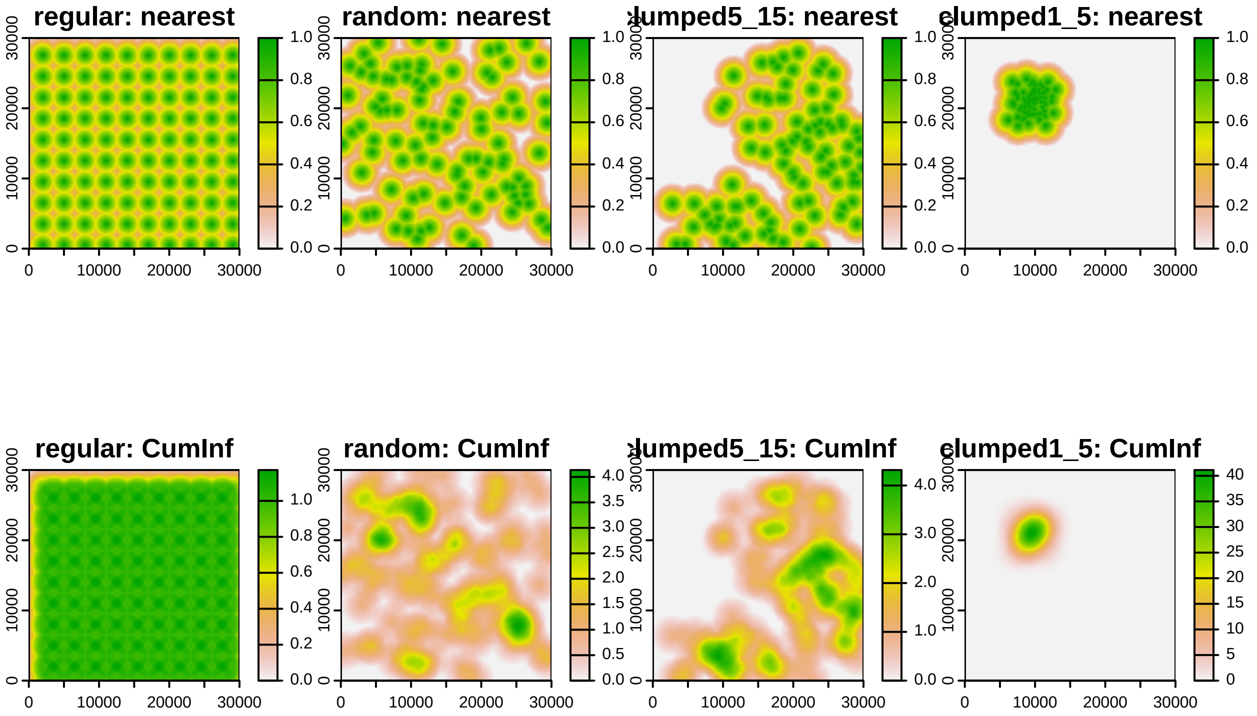

Figure C4: Illustration of the zone of influence of the nearest feature and the cumulative zone of influence of features for 4 landscapes in a gradient of clustering of the point infrastructure, for ZOI radius = 10% of the landscape extent (3000 m).

#### Correlation and variation in the ZOI metrics

To understand quantitatively how similar are $\phi_{nearest}$ and $\phi_{cumulative}$ in different conditions, we calculated the Pearson correlation coefficient between them, pixel-by-pixel. We also calculated the coefficient of variation (CV, the ratio between the standard deviation and the mean) for each measure for each landscape, as an ancillary quantity to interpret their correlation.

As seen in the plots above (Figs. C2-C4), when the minimum distance between features is larger than $2r$, the zone of influence of different features does not overlap and $\phi_{nearest}$ is similar to $\phi_{cumulative}$ (correlation = 1; Fig. C5). This happens when the ZOI of each feature is too small to overlap – when either the ZOI is small enough or there is a low number of features sparsely distributed in space. This is easily noticed for the regular distribution of points, in which the relative minimum distance between points is high (= 10% of the landscape or 3000 m; Fig. C2 and C3, Fig. C5A), but less evident in the other spatial configuration of points, since the minimum distance between points if very small (Fig. C5B-F). As $r$ increases, the two ZOI metrics diverge. Because $\phi_{nearest}$ does not account for cumulative effects, as $r$ increases its CV decreases faster than the CV of $\phi_{cumulative}$, what explains why the two metrics diverge (Fig. C6). The decrease in correlation goes on until a given ZOI radius for which the gradient represented by $\phi_{nearest}$ starts to get similar to $\phi_{cumulative}$ again[2], as shown in Fig. C5. This is mostly evident for more clustered distributions of features (Fig. C5D-F). The point of inflection is related to the radius of the clusters of features. For ZOI radii larger than the cluster radius, the variation in $\phi_{nearest}$ within the clusters vary little compared to their values outside the cluster (Fig. C6), and the cluster of features starts to act as a "super-feature" (e.g. groups of houses or wind turbines behave as urban areas or wind parks, respectively).

[2]Even though the correlation between $\phi_{nearest}$ and $\phi_{cumulative}$ increases for large ZOI radii and clustered distributions of features, the absolute scale of each ZOI metric measure is different – see the values in the color scale in Figs. C2-C4.

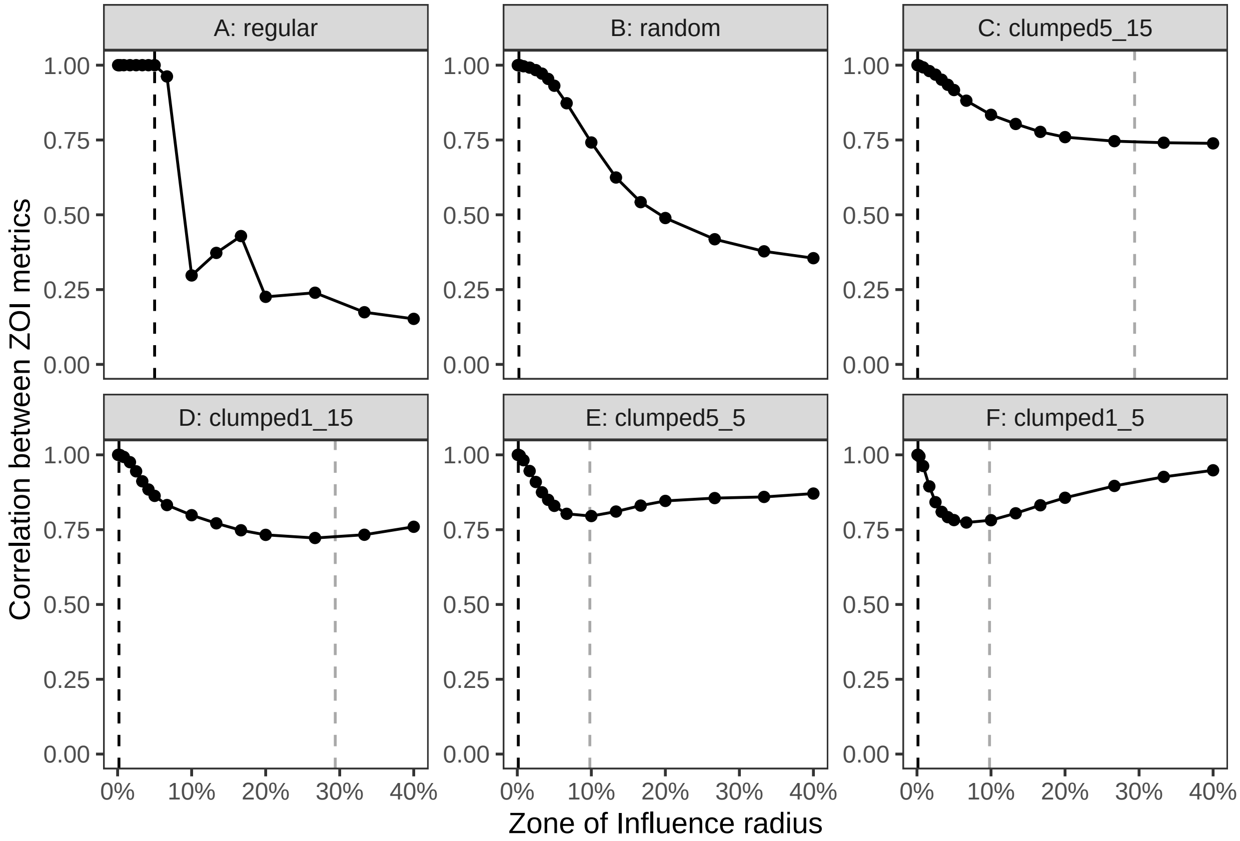

Figure C5: Pixel-to-pixel correlation between the ZOI of the nearest feature and the cumulative ZOI of multiple features as the ZOI radius increases, for the different spatial configurations of point infrastructure. The ZOI radius was varied from 0.06% to 40% of the landscape extent (i.e., from 20 m to 12 km, given our landscapes are squares of 30 km extent). The black dashed vertical line shows the minimum distance between features in a landscape, and the grey vertical dashed line shows the radius of the cluster of features, for clustered distributions.

It is important to remark that how clustered a given set of infrastructure features is depends on the extent of the study area. As an example, if the biological response is measured and assessed in a study area that comprises 2 km around a wind farm whose turbines are spread in a few mountain tops, the distribution of wind turbines might be random or somehow aggregated. However, if the study area comprises a much wider area and the biological response is expected to respond at larger extents, and the wind farm is only located in a small part of that area, their distribution will be very clumped.

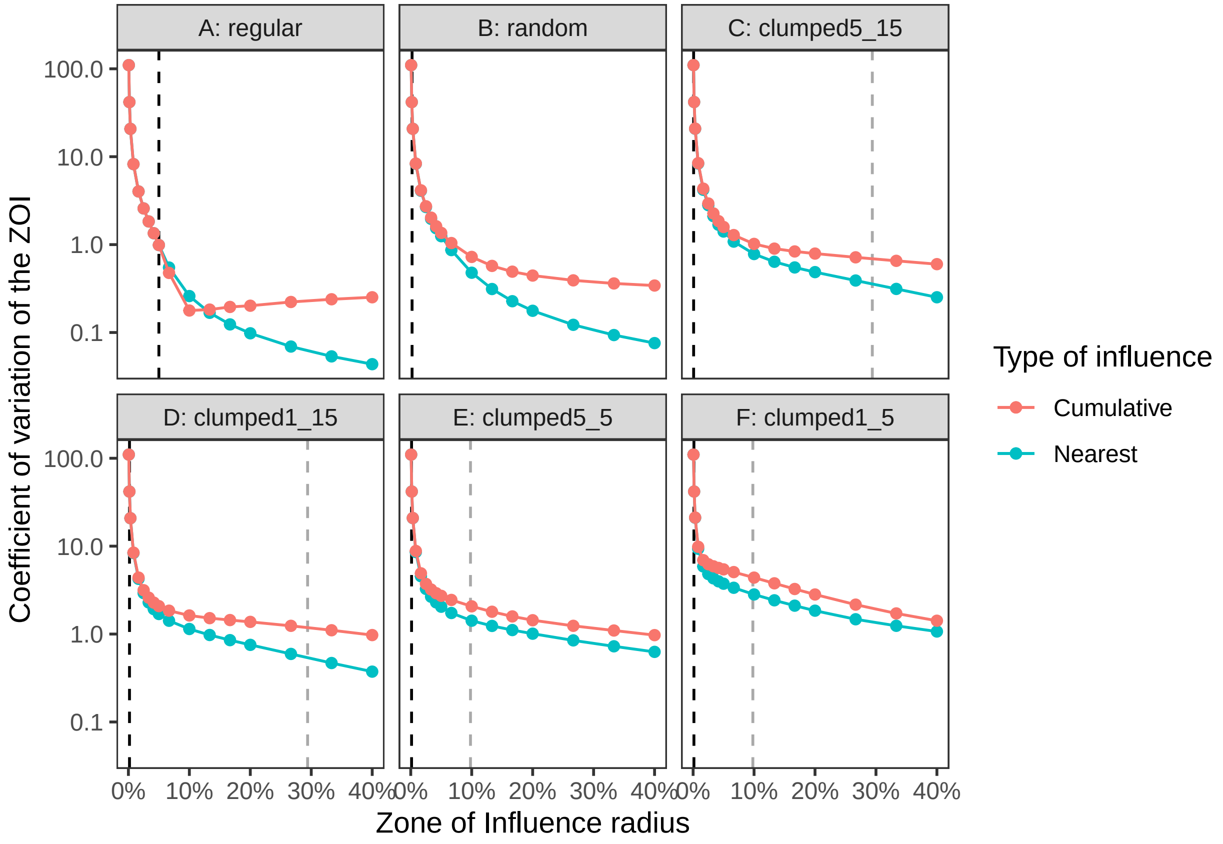

Figure C6: Coefficient of variation (CV) of the ZOI of the nearest feature and the cumulative ZOI of multiple features as the ZOI radius increases, for the different spatial configurations of point infrastructure. The ZOI radius was varied from 0.06% to 40% of the landscape extent (i.e., from 20 m to 12 km, given our landscapes are squares of 30 km extent). The y-axis is presented in log10 scale. The black dashed vertical line shows the minimum distance between features in a landscape, and the grey vertical dashed line shows the radius of the cluster of features, for clustered distributions.

### Zone of influence metrics in a real landscape

#### Context: the Hardangervidda wild reindeer area

We also provide a real-world example to illustrate the similarities of $\phi_{nearest}$ and $\phi_{cumulative}$. We use the Hardangervidda wild reindeer area in Southern Norway (Fig. 5 in the main text, Fig. D1), the region used for the habitat selection analyses presented in the main text. For more details, see the Appendix D. Similar to what was done above with the simulated landscapes, we calculated both $\phi_{nearest}$ and $\phi_{cumulative}$ for private cabins and large tourist resorts for ZOI radii varying from 100 m to 20 km, which corresponds to ~0.1 to 20% of the landscape extent (given the dimensions of the Hardangervidda reindeer area are at the order of 100 km). We calculated $\phi_{nearest}$ and $\phi_{cumulative}$ using multiple possible influence functions: linear decay, exponential decay, Gaussian decay, and threshold (see Appendix A), using the `calc_zoi()` function from the `oneimpact` R package. For each of them we calculated the pixel-to-pixel correlation and the CV, which are shown in Figs. C7 and C8.

#### Correlation and variation in the ZOI metrics

The pattern of correlations among influence measures for private cabins follow the same pattern predicted for clustered spatial points (Fig. C5), what is reasonable given their high density and aggregated spatial distribution (see Fig. D2 in Appendix D and Fig. 5 in the main text). Since the minimum distance between private cabins is very small, even for the smallest values of ZOI radius $\phi_{nearest}$ and $\phi_{cumulative}$ are already different, and for all ZOI functions their correlation decreases with ZOI radius until $r \sim 5000$ m (Fig. C7A), apart from the threshold function (for which there is no variation in $\phi_{nearest}$ for $r > 5000$ m, since all the function values are 1; see Appendix D). We did not calculate measures of or tested for clustering for these datasets, but the ZOI radius for which the correlation stops decreasing for private cabins is related to the cluster size – on relative terms, the distance between private cabins features is small (Table D1). In contrast, for public resorts the correlation among the ZOI metrics does not stop decreasing with the ZOI radius (Fig. C8A), which is also the pattern expected from the predictions from our simulated landscapes for random and sparse distributions of features (see the relative distance to the nearest tourist resort in Table C1). Here it would be necessary to go much beyond $r = 20$ km to detect some clustering in the spatial distribution of tourist resorts, a scale in which the two $\phi$ could stop diverging.

Looking at the variation in both ZOI metrics for each infrastructure type, we see that for private cabins – dense and aggregated – $\phi_{cumulative}$ is more variable than $\phi_{nearest}$ for most $r$, what explains their divergence, and their correlation stops decreasing when the CV for $\phi_{nearest}$ gets next to zero (Fig. C7B). For tourist resorts – sparse and spread in space – the CV of both ZOI metrics is similar for small $r$, and increases with $r$ (Fig. C8B).

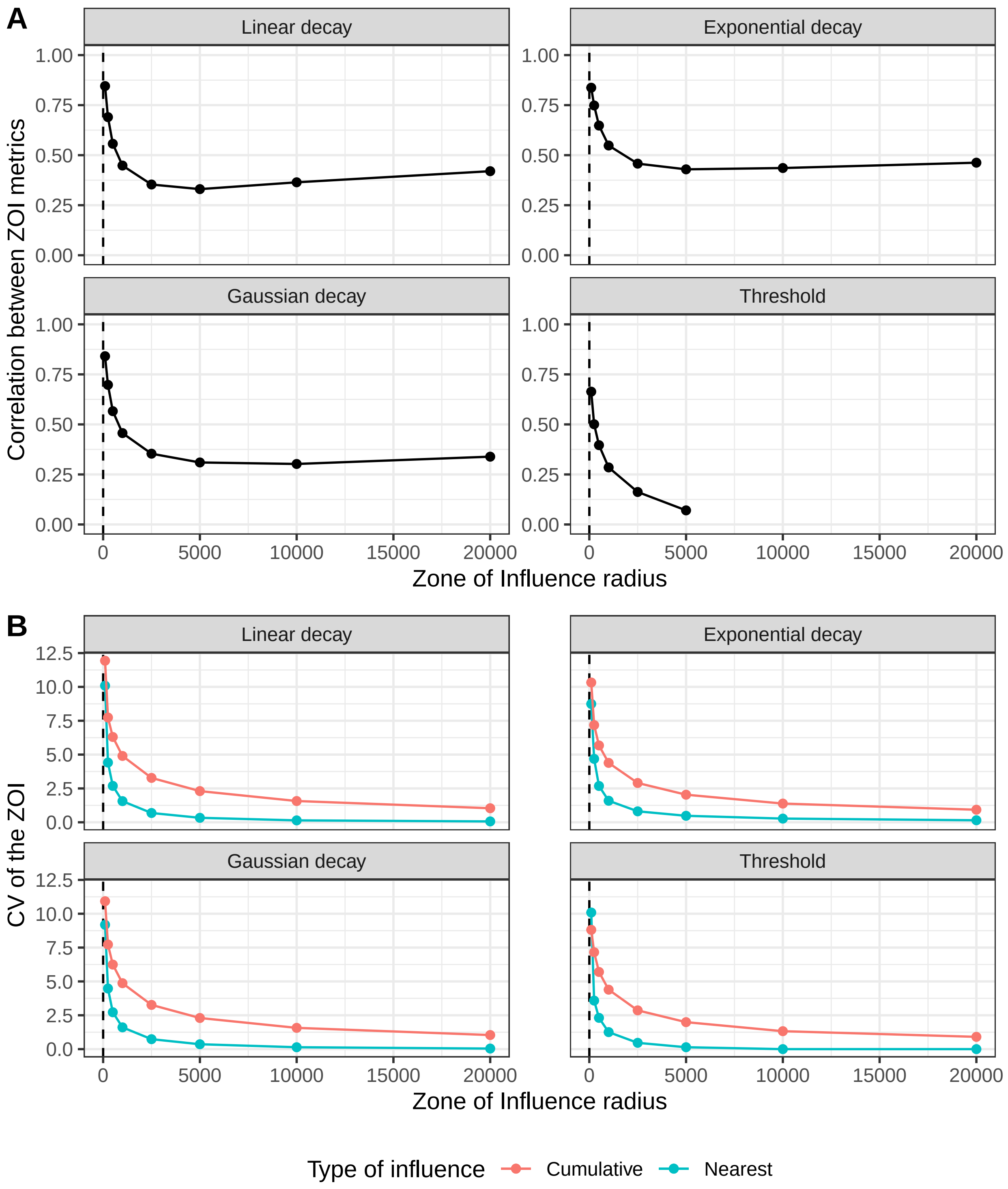

Figure C7: (A) Pixel-to-pixel correlation and (B) coefficient of variation of the ZOI of the nearest feature and the cumulative ZOI of multiple features as the ZOI radius increases, for private cabins in the Hardangervidda wild reindeer area. The vertical dashed line represents the minimum distance between features. The correlation between measures is not shown for the threshold function with $r \geq 2500m$, for which the CV of $\phi_{nearest}$ is null.

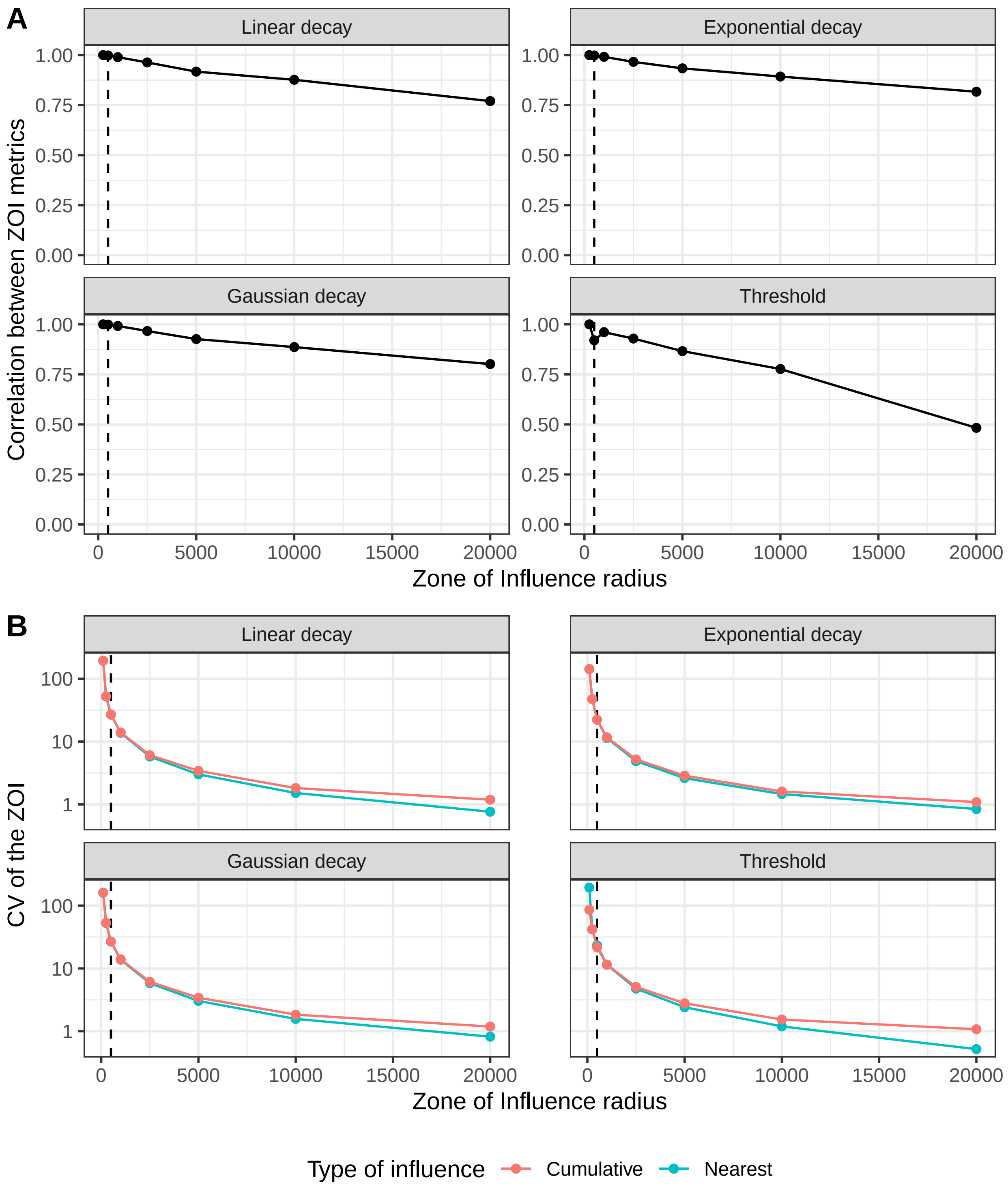

Figure C8: (A) Pixel-to-pixel correlation and (B) coefficient of variation of the ZOI of the nearest feature and the cumulative ZOI of multiple features as the ZOI radius increases, for tourist resorts in the Hardangervidda wild reindeer area. The vertical dashed line represents the minimum distance between features.

#### Additional remarks

Here we showed that the two ZOI metrics of influence – $\phi_{nearest}$ and $\phi_{cumulative}$ – are more correlated either when the spatial distribution of features is sparse and the ZOI radius is small (when the minimum distance between features is larger than the $2r$) or when features are clustered and ZOI is large ($r$ is larger than the size of clusters of features). When the correlation between $\phi_{nearest}$ and $\phi_{cumulative}$ is high it might be difficult to distinguish whether the impacts of a given infrastructure type accumulate or not. However, whether a given biological response variable is affected by only the nearest feature or the cumulative ZOI of multiple features (or none) remain an empirical problem to be explored in real landscapes.

The results presented here assume anthropogenic features represented as points, chosen here because of the simplicity to represent them and place them following different patterns. Yet, these cases also provide insights on when $\phi_{nearest}$ and $\phi_{cumulative}$ are similar for other types of infrastructure, for instance linear infrastructure (e.g. roads, railways, and power lines) and higher dimensional landscape changes defined by polygons and areas (e.g. dams, mining or forestry areas, and deforestation sites). However, we recognize that those patterns can get more complex as these structures and spatial patterns extend over large distances and areas, and a further assessment of when these ZOI metrics converge might be needed.
