## Appendix D for "Estimating the cumulative impact and zone of influence of anthropogenic features on biodiversity"

Cumulative impacts of infrastructure on reindeer space use: fitting habitat selection models

**Abstract**

In this document we fit habitat selection models to wild reindeer GPS data to assess if and how the impacts of multiple tourist infrastructure affect mountain reindeer (*Rangifer tarandus*) habitat selection during summer. We describe the modeling approach and present the results and predictions from the fitted models. This document complements the description of materials and methods and the results presented in the main text of Niebuhr et al. *Estimating the cumulative impact and zone of influence of anthropogenic features on biodiversity.*

**References** **10**

### Introduction

In this document we describe the procedures to fit habitat selection models to wild reindeer GPS data and assess if and how the impacts of multiple infrastructure affect mountain reindeer (*Rangifer tarandus*) habitat selection during summer. We first briefly describe the study area, the GPS data handling, and the environmental variables used in the analysis. We then describe the calculation of the infrastructure-related covariates using both ZOI metrics, the cumulative zone of influence (ZOI) and the ZOI of the nearest feature. These metrics quantify the ZOI as well as how the influence varies with the distance to infrastructure. Then, we describe the structure of the statistical models and the fitting procedures and present the results in details. We explore qualitatively the interpretation of both ZOI metrics in single-infrastructure models, and then estimate the effect size and the ZOI of each infrastructure in multi-infrastructure models, to finally assess the combined impacts of infrastructure on reindeer habitat selection.

### Material and Methods

#### Study area

The study area was the Hardangervidda wild reindeer area in Southern Norway, where the largest remaining population of mountain reindeer is found (Fig. D1). During summer, the area is mainly used for tourism. Hardangervidda is a big plateau surrounded by large roads around its contour, which corresponds to the lower part of the area (Fig. D2). Towards the upper, central part, there are small access roads that link the large highways to tourist resorts and a multitude of private cabins, which are also connected by a network of trails (Fig. D2). The area has 26 large tourist resorts which are constantly visited by many tourists and 24 smaller public cabins. In contrast, 14154 private cabins are spread throughout Hardangervidda. Infrastructure data was retrieved from the N50 map, obtained from GeoNorge map catalog (https://kartkatalog.geonorge.no/).

Due to their high density, most areas (90%) in Hardangervidda are closer than 3 km from any private cabin and 5 km from the closest trail (Table D1). In contrast, more than 50% of the areas are farther than 13 km from large tourist resorts and 10 km from small tourist cabins. There are also many areas far from roads towards the central part of the Hardangervidda (Table D1).

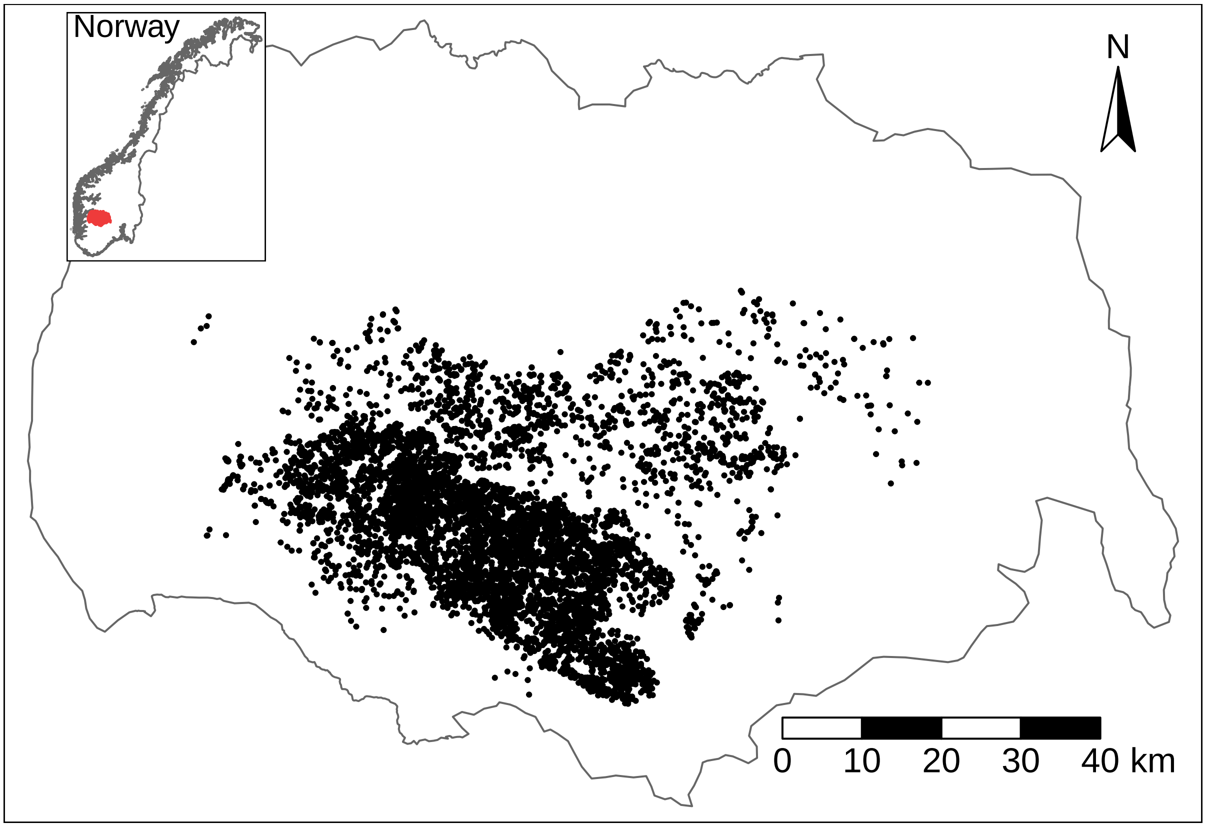

Figure D1: Hardangervidda reindeer area in Southern Norway and reindeer GPS locations for the Summer season, used in this study.

#### Reindeer GPS data

Between 2001 and 2019, 115 female reindeer were captured and monitored. Reindeer were immobilized from helicopter (see details in Evans et al., 2013) and equipped with GPS collars with drop-off system. To regularize the fix rate among collars, we used 1 reindeer position every 6 hours, summing up a total of 7478 positions for all individuals. We analyzed only the data from 1 July - 15 August, selected here as a period representative of the summer, to avoid including reindeer positions during either the end of the calving season or during rut and autumn migration. For detailed data cleaning and preparation procedures, see Panzacchi et al. (2015).

To perform habitat selection analyses, for each used GPS location we created a set of 9 locations available but not used by reindeer, spread randomly within this wild reindeer area (Fig. D1). The combination of use and available locations was annotated with environmental spatial data to assess the impacts of the different infrastructure on reindeer space use.

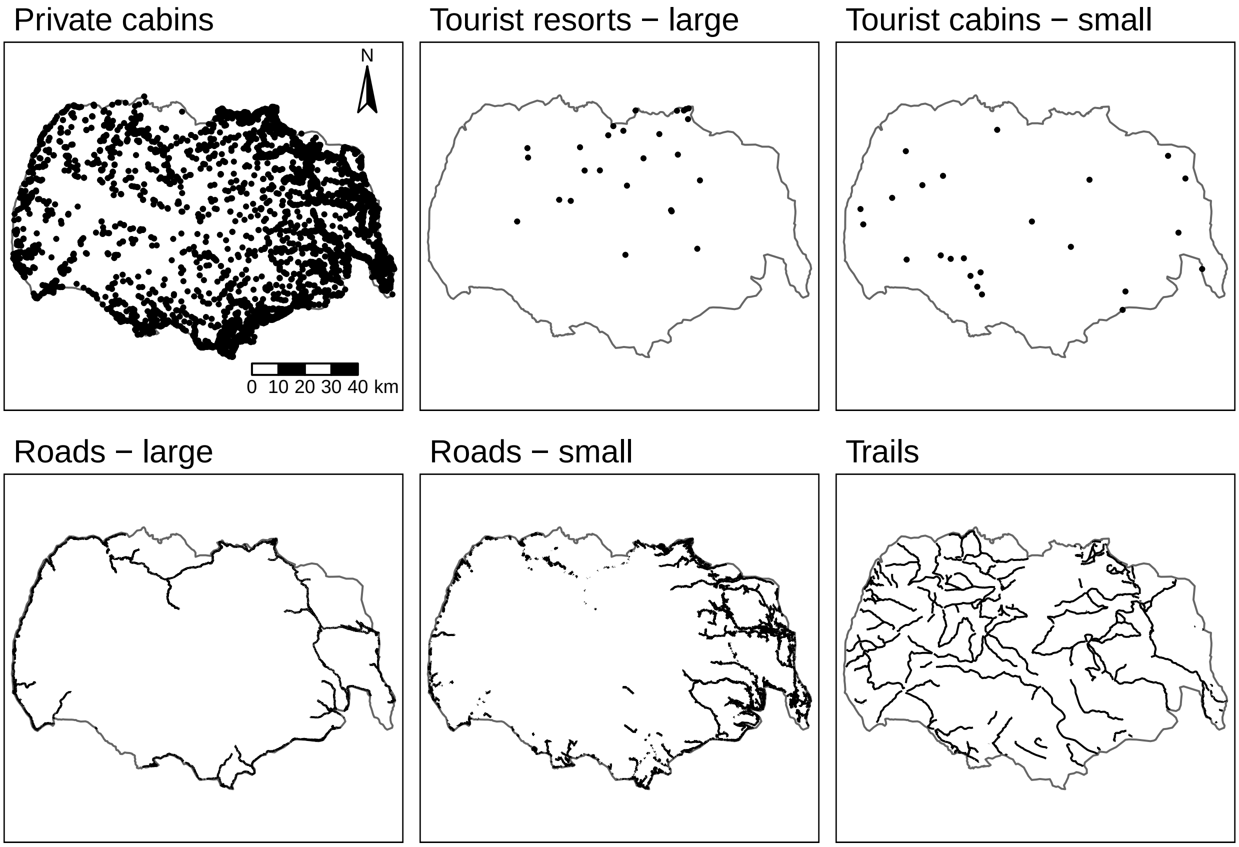

Figure D2: Main anthropogenic infrastructure in the Hardangervidda reindeer area, used to illustrate the landscape context. Only private cabins and large tourist resorts were included in the analysis.

Table D1: Quantiles of the Euclidean distance from each 100 m-side cell in the Hardangervidda reindeer area to the nearest feature (in meters), for the main anthropogenic infrastructure present in the study area.

| Infrastructure | 0% | 10% | 25% | 50% | 75% | 90% | 100% |
|---|---|---|---|---|---|---|---|
| Private cabins | 0 | 316 | 671 | 1265 | 2102 | 3178 | 7580 |
| Large tourist resorts | 0 | 4115 | 7117 | 13180 | 22496 | 29461 | 45393 |
| Small tourist cabins | 0 | 3833 | 6537 | 10308 | 14223 | 18065 | 31752 |
| Large roads | 0 | 906 | 2816 | 7235 | 14454 | 20873 | 29362 |
| Small roads | 0 | 316 | 1273 | 3970 | 9405 | 15180 | 26249 |
| Trails | 0 | 283 | 762 | 1709 | 3228 | 5124 | 11309 |

#### Environmental covariates

The locations of most types of infrastructure in Hardangervidda are correlated. Roads occur mostly in the lower parts of the area – and are correlated with elevation and terrain ruggedness – while other infrastructure occur closely together (e.g. small roads and cabins). For this reason, and for illustration purposes, in the analyses presented here we assessed only the impacts of private cabins and large tourist resorts. The spatial data sources and details are described in Panzacchi et al. (2015).

First, the vector representation for each kind of infrastructure was rasterized using a grid of 100 m resolution for an extent which included a buffer of 50 km around the study area; the buffer was used to avoid edge effects in the ZOI metrics' calculation. Then, both the ZOI of the nearest feature and the cumulative ZOI metrics were calculated. Since the infrastructure considered here (cabins and resorts) are represented as points, the input for influence calculation was the count of features within each grid cell. Rasterization was made in GRASS GIS (GRASS

Development Team, 2017) using the `v.to.rast` module and the `grass_v2rast_count` function from the `oneimpact` R package.

Influence measures were calculated considering ZOI function with different shapes (threshold, linear, Gaussian, and exponential decays; see Appendix A), for a set of (irregularly distributed) radii, from 100 m to 20 km. This allowed us to assess whether habitat selection is affected by either the cumulative impact or the impact of the nearest feature, while estimating the ZOI radius and accounting for the shape of the ZOI of the features of the different types of infrastructure. All ZOI functions were considered to have value 1 at the origin (where the infrastructure are located) and vary according to the different shapes. For the threshold and linear decay functions, the ZOI radius was defined as the distance at which the influence decreases to zero. For the Gaussian and exponential decay functions, which asymptotically approach zero, the ZOI radius was defined as the distance at which the functions reach a limit value of 0.05 (see Appendix A). The ZOI variables were computed with the `calc_zoi()` function of the `oneimpact` package using a R-GRASS GIS interface (with parameter `where = "GRASS"`). The different ZOI shapes and radii were computed by changing the parameters `type` and `radius` in this function.

To account for bio-climatic and geographic variation in reindeer space use, we also included as covariates land cover and 4 principal components (PCA axes) from a large principal component analysis performed in Norway to understand patterns of bio-climatic-geographical variation across the country (Bakkestuen et al., 2008). We used the SatVeg land cover map (Johansen, 2009) with 30 m resolution and 25 vegetation classes, which we further grouped for modeling purposes (see the final classes in Table D3). The four bio-climatic principal components represent gradients of (1) PC1 - continentality, (2) PC2 - altitude, (3) PC3 - terrain ruggedness, and (4) PC4 - solar radiation, and account for 75 - 85% of the bio-climatic variation in Norway, representing the major environmental gradients in the study area (Panzacchi et al., 2015). Prior to the analyses, the continuous variables (all but land cover) were standardized to mean 0 and standard deviation 1 using the `scale()` function in R.

#### Habitat selection modeling

Reindeer habitat selection was modeled through habitat selection functions (HSF, eq. 1 in the main text) considering the additive effect of the covariates described above. We included a quadratic term for PC1 and PC2 to account for non-linear responses (Panzacchi et al., 2015). HSFs were fitted through binomial generalized linear models using the function `glm` in R (R Core Team, 2021, with parameter `family = binomial`), with weight $w = 1$ for used locations and $w = 5000$ for available locations (as suggested in Fieberg et al., 2021).

The first step in the modeling approach was to fit HSFs considering one infrastructure type at a time in a procedure of variable selection (Burnham & Anderson, 2002), to infer which ZOI shape and radius better explained habitat selection, while also checking for correlations among the predictors (an approach similar to Laforge et al., 2015, and Huais, 2018). These models included land cover and the bio-climatic PCAs, in addition to either the cumulative ZOI or the ZOI of the nearest feature of a single infrastructure type. Given that the ZOI metrics could assume 2 representations (cumulative, nearest) and follow 4 different shapes (threshold, linear, Gaussian, exponential decay) with 8 distinct ZOI radii (100 m, 250 m, 500 m, 1 km, 2.5 km, 5 km, 10 km, 20 km), for each infrastructure type we fitted 64 HSFs. Additionally, we also fitted HSFs considering the log-distance to the nearest feature, which is a predictor commonly used in statistical models to assess the impacts of anthropogenic infrastructure on biodiversity (e.g. Torres et al., 2016; Polfus et al., 2011). Single-infrastructure HSFs were fit with the `multifit` function in R (Huais, 2018). HSFs were compared though the Akaike information criterion (AIC), and for each infrastructure type the 15 ZOI variables that better explained habitat selection (lower AIC) were chosen to be included in the multi-infrastructure HSF (see below).

We considered variables to be correlated if the Pearson correlation coefficient between their values was higher than 0.6, and excluded models in which any of the infrastructure influence measures was correlated with the bio-climatic variables.

In the single-infrastructure HSFs, we also assessed the estimated coefficients related to the infrastructure ZOI variables ($\beta$'s in Eq. 2 of the main text). Even though the coefficient values were not a criterion for selecting the most parsimonious ZOI variables, they are important to indicate consistency in the ZOI measures across the scales. If the coefficient changes signs as the ZOI radius increases, representing a shift from avoidance to selection, this might be a warning to be careful in the evaluation of the most plausible ZOI. Since the continuous covariates were standardized for model fitting, their model coefficients were rescaled back to the original covariate range, for interpretation purposes and prediction.

We fitted multi-infrastructure HSFs by combining the best ZOI variables for each infrastructure. Since not necessarily the best ZOI shape and radius for single-infrastructure models will remain as the most likely in multi-infrastructure models, we selected the 15 best covariates for each infrastructure type and fitted all possible combinations between them. For models in which the infrastructure covariates were correlated, we excluded those variables with higher Variance Inflation Factor (VIF, which measures how much the variance of an estimated regression coefficient is increased because of collinearity; Kutner, 2005). In total, we fitted $15^2 = 225$ multi-infrastructure HSFs, which were also compared through AIC.

To quantify the impacts of infrastructure, for the most likely model we used eq. 3 of the main text and multiplied the effect size – the coefficients of the fitted model – by the ZOI variable included the model. We then estimated habitat suitability by predicting the HSF (eq. 1 in the main text) over the space and rescaling the predicted values to the interval [0, 1].

### Results

#### Single-infrastructure HSF

We start by describing how much support the different ZOI variables presented in explaining reindeer habitat selection in the single-infrastructure models. By doing so, we aim at showing qualitatively what the different influence measures represent and how one would interpret them within an ecological context.

##### Private cabins

For private cabins, the most parsimonious HSF included the cumulative ZOI with Gaussian decay shape and radius $r = 10$ km, but the support for the cumulative ZOI with the same radius of 10 km but other shapes was also relatively high (low relative difference in AIC, Fig. D3A). Overall, the models including cumulative ZOI metrics (regardless of the ZOI function shape and in great part of the ZOI radius) performed much better than the ones including the ZOI of the nearest private cabin (Fig. D3A), what points to strong evidence that the impacts of multiple private cabins on reindeer habitat selection accumulate. The coefficients were consistently negative across ZOI radii (Fig. D3B), which indicates the ZOI radii with minimum AIC presented in the x axis of the Fig. D3A are also consistent.

We also go beyond the simple statistical variable selection and interpret the most parsimonious models considering the ZOI of the nearest feature. In this case, regardless of the ZOI shape, the ZOI radius varied from 500 m to 1000 m (Fig. D3A). Combining the results, we can say that, if we consider the closest private cabin only, reindeer generally avoid being closer than 1 km from any cabin, but since many areas have a high density of cabins (Fig. D2, Table D1), they respond to the combined impact of many individual cabins at a larger extent - a zone of influence of 10 km radius. This might also be related to how the cabins are used. Tourists who stay in a cabin hardly walk farther than a few kilometers from it, since they must return to the cabin in the end of the day. Then, the ZOI radius of a single cabin is shorter. However, in areas where many private cabins are clustered, there is a much wider area used by tourists and the radius of the ZOI of this combined cluster of cabins is higher.

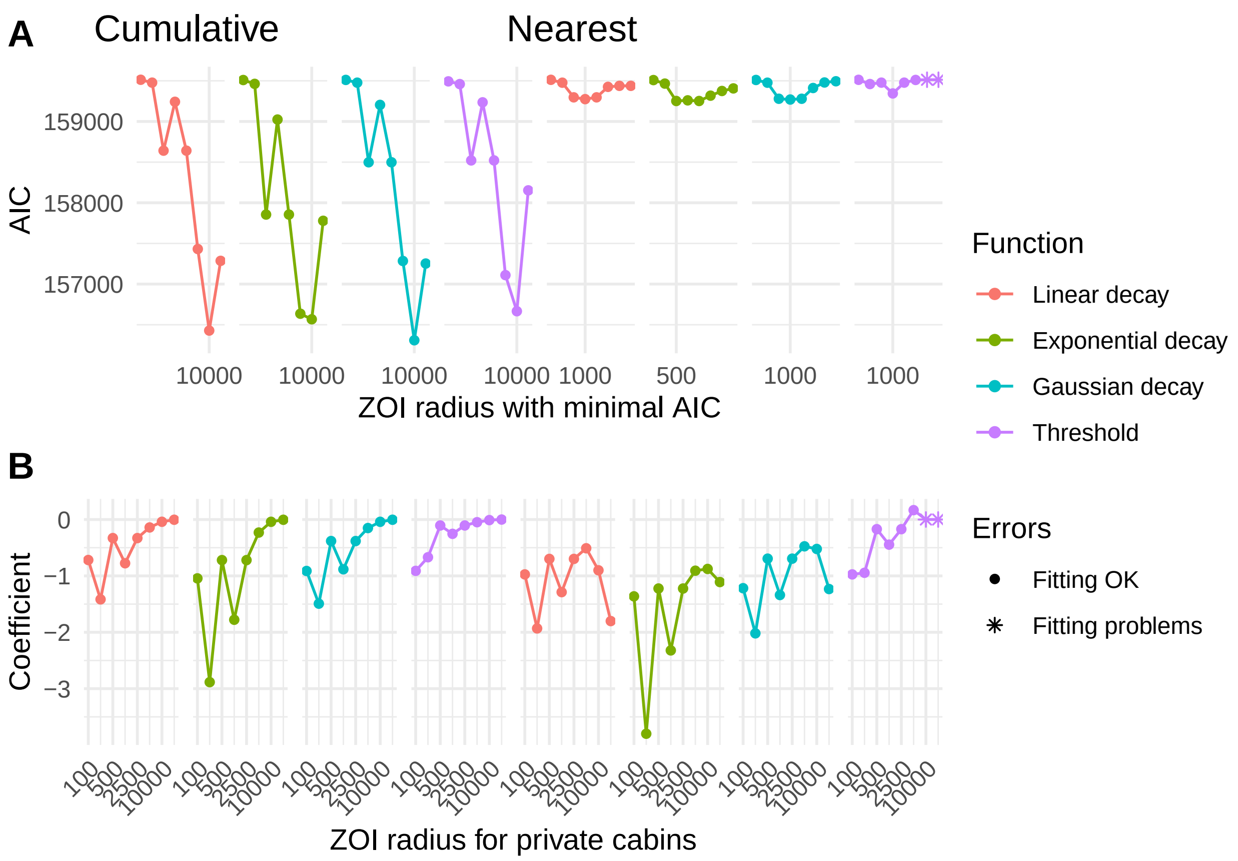

Figure D3: Model AIC (A) and coefficients (B) estimated for the zone of influence of private cabins in models including only this type of infrastructure. The plots show the AIC and the coefficients (scaled back to the original range of the covariates) of the cumulative ZOI and the ZOI of the nearest feature for different ZOI shapes and radii (see the x axis in B for some of the candidate ZOI radii values, in meters, which varied from 100 m to 20 km). The x axis in A shows the ZOI radius at which the AIC was minimal for each ZOI metric and shape. Scales marked with '*' represent errors in the model fitting (e.g. threshold ZOI of the nearest feature for ZOI = 10 km or 20 km, when the variable is constant = 1 over the whole study area).

**Large tourist resorts**

In the single-infrastructure models, there was also evidence of cumulative impacts of large tourist resorts with a 20 km zone of influence (Fig. D4). The most supported ZOI variable was the cumulative ZOI with exponential decay shape and radius of 20 km. This ZOI radius was selected regardless of the function shape, and even for the ZOI of the nearest feature the most common radius was 20 km (see the x axis in Fig. D4A).

The closer correspondence between the selected radius of cumulative ZOI and nearest ZOI metrics, in comparison to the private cabins, might be due to several factors. First, tourist resorts are present at a much lower density in Hardangervidda, with high median distance to the closest resort (Fig. D2 and Table D1). As a consequence, there is not so much difference between what the cumulative and nearest ZOI metrics represent (as for the private cabins), since the ZOI of each feature only start to accumulate for larger radii. Therefore, the ZOI corresponding to smaller AIC is closer between models including cumulative and nearest metrics. This is in accordance with the demonstrations from Appendix C, where we showed that, for landscapes with sparse distribution of features and small ZOI radius, $\phi_{nearest}$ and $\phi_{cumulative}$ are highly correlated. Second, large tourist resorts are used very differently by tourists than private cabins. Each resort is visited by a large load of tourists at a time, which consequently use a much wider area around the resorts, compared to each single private cabin. As a consequence, the ZOI radius of large tourist resorts tend to be higher.

As for the private cabins, in the models with tourist resorts the coefficients were also consistently negative, representing reindeer avoidance to public tourist resorts (Fig. D4B).

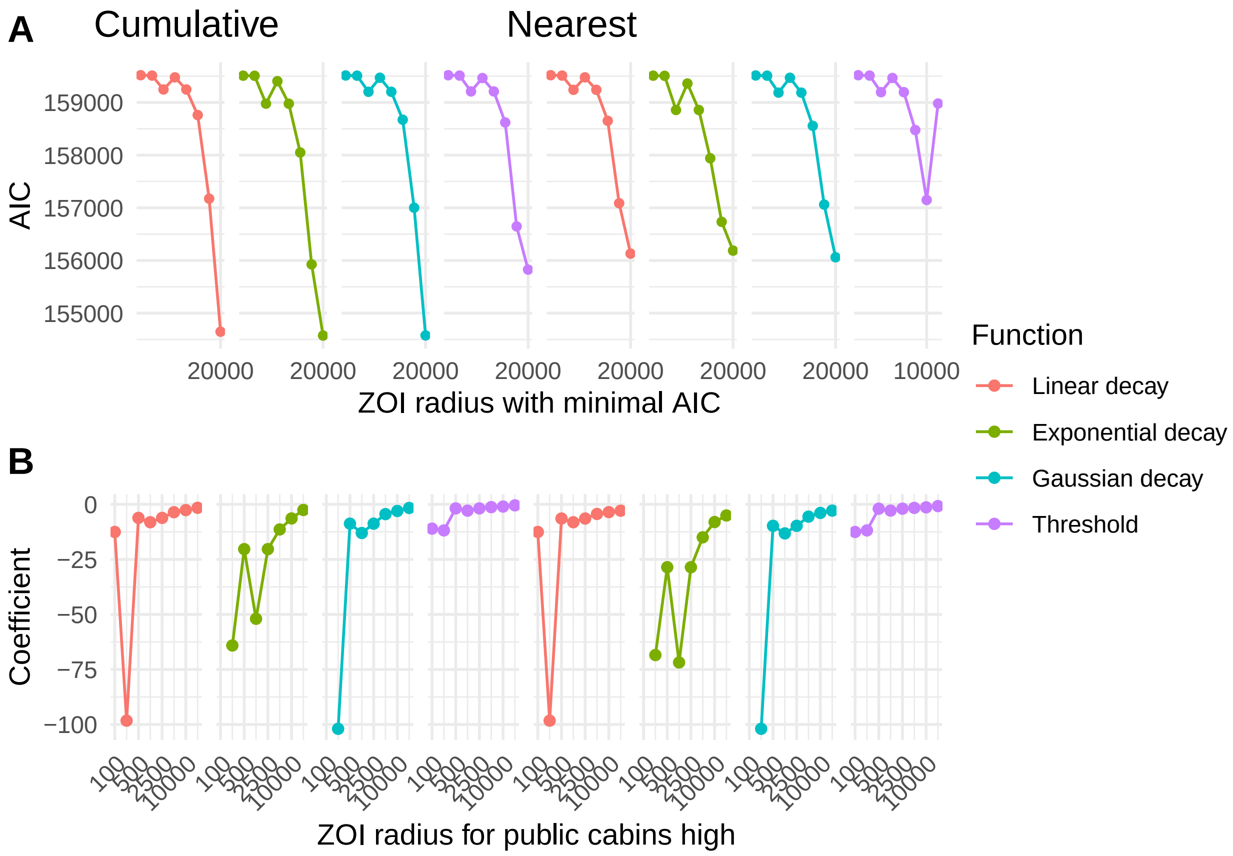

Figure D4: Model AIC (A) and coefficients (B) estimated for the zone of influence of large tourist resorts in models including only this type of infrastructure. The plots show the AIC and the coefficients (scaled back to the original range of the covariates) of the cumulative ZOI and ZOI of the nearest feature for different ZOI shapes and radii (see the x axis in B for some of the candidate values for the radius, in meters, which varied from 100 m to 20 km). The x axis in A shows the ZOI radius at which the AIC was minimal for each ZOI metrics and shape.

#### Multi-infrastructure HSF

The most parsimonious multi-infrastructure model included the cumulative ZOI of private cabins with threshold decay and $r = 10$ km and the cumulative ZOI of multiple tourist resorts with exponential decay and $r = 20$ km ($\Delta AIC = 26.9$ from the second-ranked model, $wAIC = 1$; Table D2). Notice that, as parameterized here, for the tourist resorts an exponential decay ZOI with radius of 20 km means that the influence of resorts decrease to half of its maximum value at ca. 5 km from the infrastructure (exponential half life is $\sim ZOI/4$ here). The most plausible model with a covariate for the ZOI of the nearest feature was ranked 26$^{\text{th}}$ in the model selection ($\Delta AIC = 921$), and the most likely model including the log-distance to the nearest feature was ranked 44$^{\text{th}}$ ($\Delta AIC = 1197$; Table D2). This presents strong support for the cumulative impacts of both private cabins and tourist resorts on reindeer habitat selection in Hardangervidda.

Table D2: Infrastructure variables included in the most parsimonious models. Models ranked after the 5th place are omitted. For each model we show the type of ZOI metric ("cumulative", "nearest"), the ZOI function ("exponential decay", "gaussian decay", "threshold", "Bartlett or linear decay"), and the ZOI radius (in km) for that covariate included in the model. For each model we also present the number of parameters k, AIC, the difference in AIC to the most likely model (dAIC), and the AIC weight. The last lines show the most plausible model which included any variable with the ZOI of the nearest feature and the log-distance to the nearest feature (in this case, for tourist resorts). Models also included bio-climatic variables and land cover (see Table D3).

| Rank | Private cabins | Large tourist cabins | k | AIC | dAIC | wAIC |
|---|---|---|---|---|---|---|
| 1 | cumulative, threshold, 10 | cumulative, exp decay, 20 | 23 | 152167 | 0 | 1 |
| 2 | cumulative, exp decay, 10 | cumulative, exp decay, 20 | 23 | 152194 | 26.9 | <0.001 |
| 3 | cumulative, Gauss, 10 | cumulative, exp decay, 20 | 23 | 152212 | 45.7 | <0.001 |
| 4 | cumulative, exp decay, 20 | cumulative, exp decay, 20 | 23 | 152247 | 80.1 | <0.001 |
| 5 | cumulative, Gauss, 20 | cumulative, exp decay, 20 | 23 | 152280 | 112.7 | <0.001 |
| 26 | cumulative, Gauss, 20 | nearest, bartlett, 20 | 23 | 153088 | 921.3 | <0.001 |
| 44 | cumulative, exp decay, 10 | nearest, log nearest, NA | 23 | 153364 | 1197.4 | <0.001 |

Looking closely to the most plausible multi-infrastructure HSF (after rescaling the coefficients back to the original range of variation of the infrastructure ZOI predictors), we see both anthropogenic feature types are avoided by reindeer. Their impact vary differently across space since their ZOI functions and radii differ – a threshold ZOI with 10 km radius for private cabins and an exponential decay ZOI with 20 km radius for tourist resorts –, but also because the estimated effect size of a single private cabin ($\beta_{\text{private cabin}} = -0.0081$) was much smaller than that of a single tourist resort ($\beta_{\text{tourist resort}} = -2.654$; Table D3, Fig. D5A). However, since private cabins occur at much higher densities, in some areas their overall impact is higher than that of tourist resorts (Fig. D5, Fig. 4 in the main text). Comparing an area with only 1 cabin in a 10 km radius with an area with only 1 tourist resorts in a 20 km radius, and assuming all other conditions are similar, the impact – measured here as the product between the effect size and the ZOI covariate (eq. 5 from the main text) – is much smaller for private cabins (Fig. D5A). In contrast, if we take the areas with higher influence in Hardangervidda – where the number of private cabins sum to 2664 and the (exponentially weighted) number of tourist resorts sum to 5 – the impact of private cabins agglomerates is higher than that of tourist resorts (Fig. D5C). Following the HSF coefficient interpretation from Fieberg et al. (2021), and considering that all other conditions are kept similar, a reindeer avoids an area 14.43 ($exp(330 \cdot 0.0081) = 14.43$) times more strongly than another area with 330 less private cabins in a radius of 10 km. That is approximately the same difference in avoidance a reindeer presents among two areas that differ in 1 tourist resort in a radius of 20 km ($exp(1 \cdot 2.654) = 14.21$).

Table D3: Effect size (model coefficients) of the most parsimonious model of space use for reindeer, including private cabins and tourist resorts. The table shows the coefficient estimates (scaled back to the scale of variation of the original data), their standard error (SE, in the standardized scale of the variables), and the significance (p). "pc" are the bio-climatic principal components and "poly" are the coefficients of a quadratic function of pc.

| Covariate | Estimate | SE | p |
|---|---|---|---|
| (Intercept) | -15.3081 | 0.16 | < 0.0001 |
| private cabins (cumulative, threshold, 10km) | -0.0081 | 0.13 | < 0.0001 |
| tourist resorts (cumulative, exponential, 20km) | -2.65438 | 0.04 | < 0.0001 |
| exposed ridges | 0.20948 | 0.14 | 0.1343 |
| grass ridges | 0.99185 | 0.13 | < 0.0001 |
| heather ridges | 0.99219 | 0.13 | < 0.0001 |
| lichen | 1.22547 | 0.17 | < 0.0001 |
| heather | 1.04047 | 0.13 | < 0.0001 |
| heathland | 0.93174 | 0.13 | < 0.0001 |
| meadows | 0.97105 | 0.15 | < 0.0001 |
| early snowbed | 0.61526 | 0.13 | < 0.0001 |
| late snowbed | 0.43487 | 0.13 | 0.0012 |
| bog | 0.97646 | 0.15 | < 0.0001 |
| glacier | -0.43692 | 0.32 | 0.1732 |
| other | -3.07193 | 37.62 | 0.9349 |
| water | -1.6334 | 0.2 | < 0.0001 |
| poly(pc1, 2)1 | 323.5927 | 10.23 | < 0.0001 |
| poly(pc1, 2)2 | -253.8435 | 10.81 | < 0.0001 |
| poly(pc2, 2)1 | -611.40893 | 38.54 | < 0.0001 |
| poly(pc2, 2)2 | -204.9332 | 22.97 | < 0.0001 |
| pc3 | 27.73861 | 23.91 | 0.246 |
| pc4 | -77.24946 | 24.02 | 0.0013 |

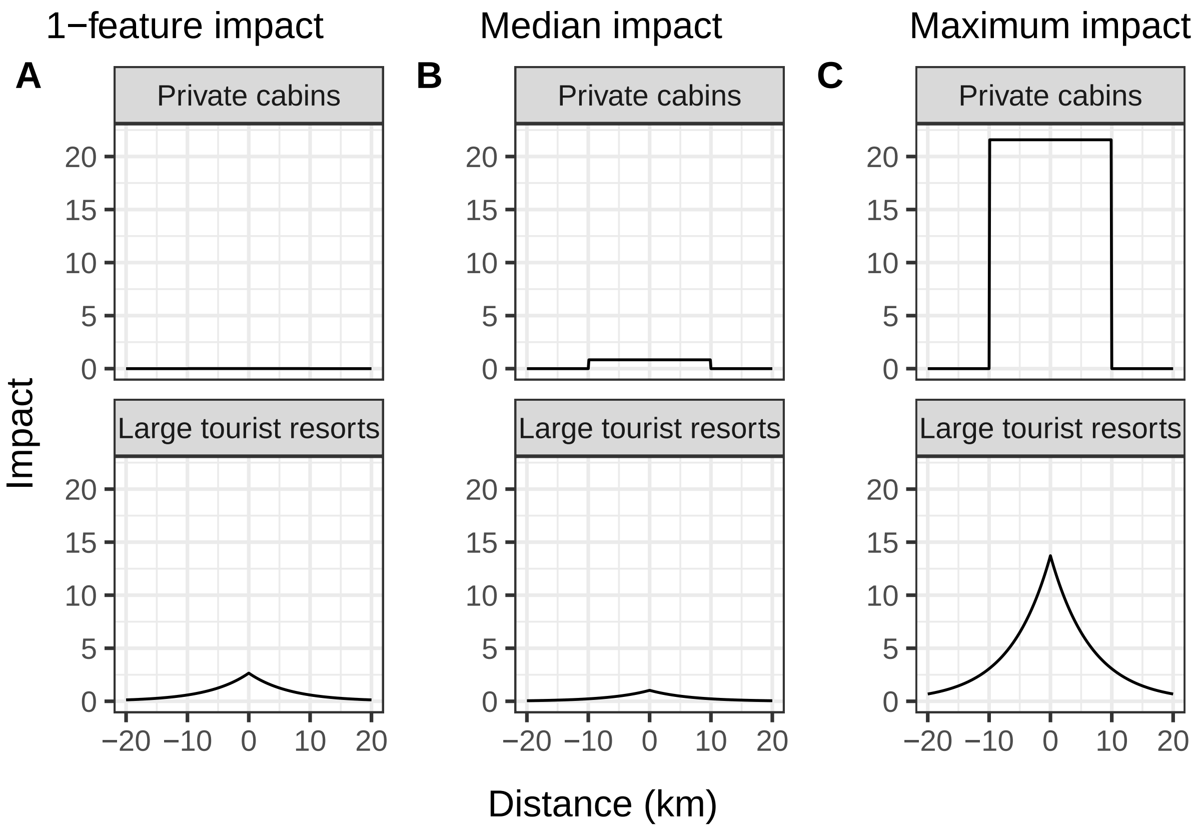

Figure D5: Impact of private cabins and tourist resorts considering (A) only 1 feature, (B) the median number of features (103 for private cabins, 0.38 for tourist resorts), and (C) the maximum number of each type of feature (2664 for cabins, 5 for resorts), given their respective estimated ZOI shape and radius The impact presented here is the multiplication between the effect size (the model coefficients) and the cumulative ZOI variable (eq. 5 in the main text). The impact of only one private cabin is negligible (A). At their median values, the impacts of private cabins and public resorts are comparable (B), while at their maximum the cumulative impact of private cabins might be higher than that of tourist resorts (C).

When cumulative impacts of infrastructure are predicted in space by multiplying the effect size and the cumulative ZOI metrics, we see how the relative impact of private cabins and large tourist resorts change across space (see Fig. 5 in the main text). Since reindeer avoided high densities of both infrastructure types at relatively large extents, areas of high habitat suitability for reindeer corresponded to those in which the cumulative ZOI of both infrastructure is low – what matches with the locations used by reindeer, indicated through the GPS data.
